# Story of an infection: viral dynamics and host responses in the *Caenorhabditis elegans*-Orsay virus pathosystem

**DOI:** 10.1101/2023.10.31.564947

**Authors:** Victoria G. Castiglioni, Maria J. Olmo-Uceda, Ana Villena-Giménez, Juan C. Muñoz-Sánchez, Santiago F. Elena

**Author notes:** These authors contributed equally to this work (Victoria G. Castiglioni, Maria J. Olmo-Uceda).

## Abstract

Orsay virus (OrV) is the only known natural virus affecting *Caenorhabditis elegans*, with minimal impact on the worm’s fitness due to its robust innate immune response. This study aimed to understand the interactions between *C. elegans* and OrV by tracking the infection’s progression during larval development. Four distinct stages of infection were identified based on viral load, with a peak in capsid-encoding RNA2 coinciding with the first signs of viral egression. Transcriptomic analysis revealed temporal changes in gene expression and functions induced by the infection. A specific set of up- regulated genes remained active throughout the infection, and genes correlated and anticorrelated with virus accumulation were identified. Responses to OrV mirrored reactions to other biotic stressors, distinguishing between virus-specific responses and broader immune responses. Additionally, mutants of early response genes and defense-related processes showed altered viral load progression, uncovering new players in the antiviral defense response.

## INTRODUCTION

Viruses, the most abundant biological entities on Earth, infect all known living organisms (Suttle, 2007; Knipe and Howley, 2013). From bacteria to eukaryotes, most organisms possess immune systems to defend themselves against viruses. Viruses, in turn, adapt to their hosts by evolving strategies to evade or neutralize the host’s defenses. This ongoing arms race between viruses and their hosts has driven the development of complex interactions and defense mechanisms (Morgan and Koskella, 2011).

The nematode *Caenorhabditis elegans* MAUPAS lacks adaptive immunity (Engelmann and Pujol, 2010). In turn, it uses the innate immune system to defend itself against viruses and other microbes, such as its natural pathogen Orsay virus (OrV) (Félix et al., 2011). OrV has a positive-sense, bi-segmented RNA genome of ∼6.3 kb which encodes four proteins: an RNA-dependent RNA polymerase (RdRP), a viral capsid protein (CP), protein (δ), and a protein fusion of CP-δ (Félix et al., 2011; Jiang et al., 2014). Free δ mediates nonlytic viral release (Yuan et al., 2018) and the CP-δ fusion is incorporated into the virions as pentameric head fibers (Fan et al., 2017; Guo et al., 2020).

OrV replicates in the intestinal epithelia of *C. elegans* (Félix et al., 2011; Franz et al., 2014), which consists of 20 single-layer cells surrounding the central lumen. Infection of OrV is restricted to the anterior intestinal cells and after cell egression results in a orofecal route of transmission (Félix et al., 2011). The infection results in enlarged intestinal lumens and subcellular structural changes, as well as reduced food consumption and smaller body size. However, the infection does not affect animal life span or brood size, suggesting that the innate immune system is able to effectively combat the virus (Ashe et al., 2013).

Several conserved antiviral defense mechanisms that *C. elegans* uses against OrV infection have been described, including (*i*) the RNA interference pathway, which degrades viral RNA and triggers antiviral gene expression (Ashe et al., 2013; Chen et al., 2017; Sowa et al., 2020); (*ii*) viral RNA uridylation, which targets viral RNA to degradation (Le Pen et al., 2018); and (*iii*) expression of antiviral genes, which include protein ubiquitination for targeted degradation and the activation of the intracellular pathogen response (IPR), which is a common response to intracellular pathogens and stresses (Bakowski et al., 2014; Chen et al., 2017; Reddy et al., 2017, 2019). In addition, several studies have described pathways necessary for OrV replication, such as zinc and the lipid metabolism (Casorla-Perez et al., 2022a), chromatin remodeling and cytoskeleton rearrangements (Zhou et al., 2023).

Studies of OrV infection have allowed us to better understand virus-host interaction mechanisms. However, there is little known on the progression of the viral infection and in the dynamic interaction between *C. elegans* development and OrV during the course of an infection. In order to shed light in these matters, we have analyzed the progression of the infection and the host response in a time resolved manner in the mildly-resistant wild-type nematode strain N2. By examining the progression of the viral load, we observed that the infection could be divided into four stages: (1) pre-infection, (2) exponential viral replication leading to a viral load peak, (3) host-pathogen conflict leading to a second lower viral load peak, and (4) persistent residual infection. We noted that the ratio of the two genomic RNAs varied during the different phases of the infection, with a higher production of the RNA2 encoding capsid proteins coinciding with the peaks in viral load. Relatedly, we started seeing signs of viral egression after the first peak, suggesting that a viral generation would be complete. Remarkably, the number of infected cells and the relative infected area remained mostly stable along the infection. In order to investigate the response elicited in the worm, we looked at the changes in the transcriptome along the four infection stages. As a result, we obtained a transcriptomic landscape of the genes and functions involved in the response to infection during the worm’s development. In-depth analyses allowed us to pinpoint a set of genes up-regulated throughout the infection process and to highlight genes most correlated and anticorrelated with virus accumulation. We also saw responses to OrV that are frequently linked to other biotic stressors, suggesting that the immune response triggered by OrV may be more general than previously thought, and a unique response to OrV which differed from the response to other biotic stresses. Finally, we identified early responding genes and described novel pro-viral and anti-viral genes.

## RESULTS

### Dynamic viral accumulation during larval development shows a damped wave-like pattern

Wild-type (N2) worms are mildly-resistant to OrV infection in comparison to the JU1580 natural isolate in which OrV was found (Félix et al., 2011). JU1580 carries a mutation in the viral recognition protein DRH-1, homolog of RIG-I, which renders worms RNAi deficient (Ashe et al., 2013). However, inoculations with high doses of OrV have been previously shown to produce comparable amount of virus in N2 and *drh-1(ok3495)* (Sterken et al., 2014). Here we used a highly concentrated stock and observed similar levels of OrV in the N2-background strain ERT54 (Bakowski et al., 2014) and JU1580 worms (Mann Whitney test, *P* = 0.0556) (Fig. S1). The use of this highly concentrated stock allowed the study of a synchronous infection in wild-type populations during larval development.

We first followed the progression of the viral load in a synchronized population from hatching to the end of larval development (L4). For this purpose, we inoculated worms at the time of hatching and took samples at increasing hours post-inoculation (hpi) for quantitative RT-PCR (RT-qPCR), RNA-seq and smiFISH analyses (Fig. 1A). Viral loads analyzed by RT-qPCRs targeted RNA2 and were determined using the standard curve method. Viral fragments counted by RNA-seq were relativized to the total reads in each sample. Both analysis indicated that the progression of the viral load could be divided in four phases (Fig. 1B): (1) a pre-replication phase in which the viral load is indistinguishable from background levels (0 - 6 hpi), (2) an exponential viral replication leading to a viral load peak (8 - 12 hpi), (3) a phase of host-pathogen conflict leading to a second lower viral load peak (14 - 36 hpi), and (4) a phase of persistent residual infection in which the viral load came close to the initial background levels (38 - 44 hpi). RT-qPCR and RNA-seq quantifications of OrV showed a high correlation (*rs* = 0.9720, 10 d.f., *P* < 0.0001), confirming that both methods are reliable estimates of viral load.

**Figure 1.**
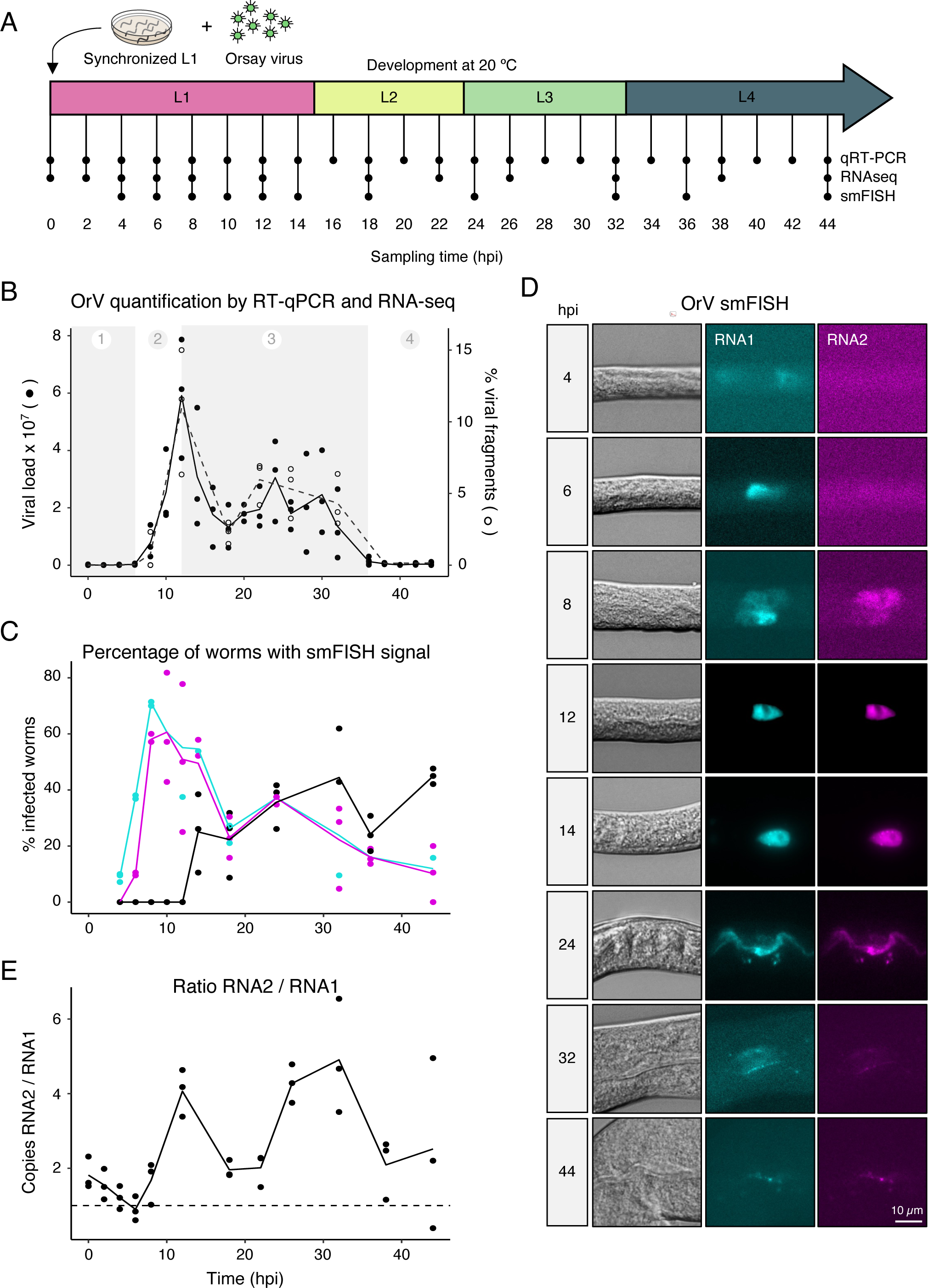
Dynamic viral accumulation during larval development (A) Experimental design. Synchronized recently hatched N2 populations where inoculated with 2.8×10^9^ copies of OrV and grown at 20 °C. Samples were analyzed at the indicated timepoints by qRT-PCR, RNA-seq or smiFISH. (B) Viral load determined by RT-qPCR (left axis, solid circles and solid line) and percentage of viral reads from the total RNA determined by RNA-seq (right axis, empty circles and dashed line) over time. The viral load by RT-qPCR was defined as copies of RNA2/10 ng of total RNA. Lines represent the mean; *n* = 3 replicates per time point. The different stages of the infection, in grey shading, are described in the main text. (C) Percentage of worms with cellular RNA1 (cyan), RNA2 (magenta) or with luminal expression of either RNA1 or RNA2 (black) over time as determined by smiFISH. Lines represent the mean; *n* = 3 replicates per time point. (D) Representative smiFISH images of inoculated worms at the indicated timepoints. Left: DIC; middle: RNA1 in cyan; right: RNA2 in magenta. (E) Ratio of RNA1/RNA2 over time previous relativization to fragment length of each genome determined by RNA-seq; *n* = 3 replicates per time point. See also Fig. S1.

Furthermore, we sought to characterize the percentage of infected worms throughout larval development. We inoculated worms at the time of hatching and took sequential samples for smiFISH analysis (Parker et al., 2021). RNA1 and RNA2 were labelled independently with 22 and 18 smiFISH probes, respectively. We observed that all the worms that expressed RNA2 also expressed RNA1, but the reverse was not necessarily true. RNA1 encodes the RdRP, therefore it would be reasonable that its levels would rise faster than RNA2 during early infection. Indeed accumulation of RdRP has been shown to precede the accumulation of the capsid proteins (Franz et al., 2014). We observed that the incidence of infection in the worm population recapitulated the same four phases described above: (1) a pre-replication phase in which the percentage of infected worms is close to zero and the smiFISH signal was hardly above background (4 hpi), (2) an exponential increase in the percentage of infected worms in which the intensity of signal quickly increased leading to a peak in infection (6 - 14 hpi), (3) a phase of host-pathogen conflict leading to a second and lower peak in which the intensity of signal was significantly reduced (18 - 32 hpi), and (4) a phase of persistent residual infection in which the percentage of infected worms was drastically reduced and the intensity of signal was again hardly above background (36 - 44 hpi) (Fig. 1C). Furthermore, the highest percentage of infected worms was observed earlier than the viral load peak, at 8 and 12 hpi respectively, which may suggest that some worms are able to resolve the infection quickly.

We next asked whether the accumulation of RNA1 and RNA2 undergoes a shift in gene expression from early to late replication cycle stages. We hypothesized that RdRP expression from RNA1 would be higher at the beginning of the viral replication cycle when viral replication is higher, while levels of structural proteins from RNA2 would became more abundant towards the end of the viral replication cycle as virions are being assembled (Félix et al., 2011). For this purpose, we looked at both RNA1 and RNA2 labelling with smiFISH and at viral counts relativized to fragment size from RNA-seq. At the initial phases of the infection a higher percentage of worms expressed RNA1 in comparison with RNA2, indicating that higher RNA1 levels are needed before transcription of RNA2 (Fig. 1D). Consistently, RNA-seq data indicated that the highest RNA1 to RNA2 ratio is at the beginning of the infection. However, RNA2 was always more abundant than RNA1, with an average 2.40 ±1.62 RNA2 to RNA1 ratio throughout all timepoints (Fig. 1E). Interestingly, the ratio shows a dynamic behavior, with higher values coinciding with the viral load peaks and the highest ratio coinciding with the second viral peak, indicating that RNA2 is more abundant before the end of the viral replication cycle and consistent with the need of RNA2-encoded products for encapsidation (Fig. 1E).

### Spatial progression of infection

Next, we characterized the spatial distribution of viral RNAs in the worm. In the initial infection phases, viral particles were found only inside intestinal cells (Fig. 1C-D, Fig. 2A). However, at 14 hpi, viral RNA was first seen in the intestinal lumen, indicating a productive release of viral particles. By 24 hpi, most worms showed a strong signal in the lumen (Fig. 2B). These findings suggest that RNA1 is prioritized for translation and transcription after viral entry. Around 6 hours post RdRP translation (Franz et al., 2014), RNA2 levels begin to rise, reaching their peak at 12 hpi. Following this peak, we notice a decline in RNA2 levels, along with a significant relocation of OrV to the lumen, signifying egression (Fig. 2C). Overall, these results imply that a viral replication cycle might conclude in as little as 14 hpi.

**Figure 2.**
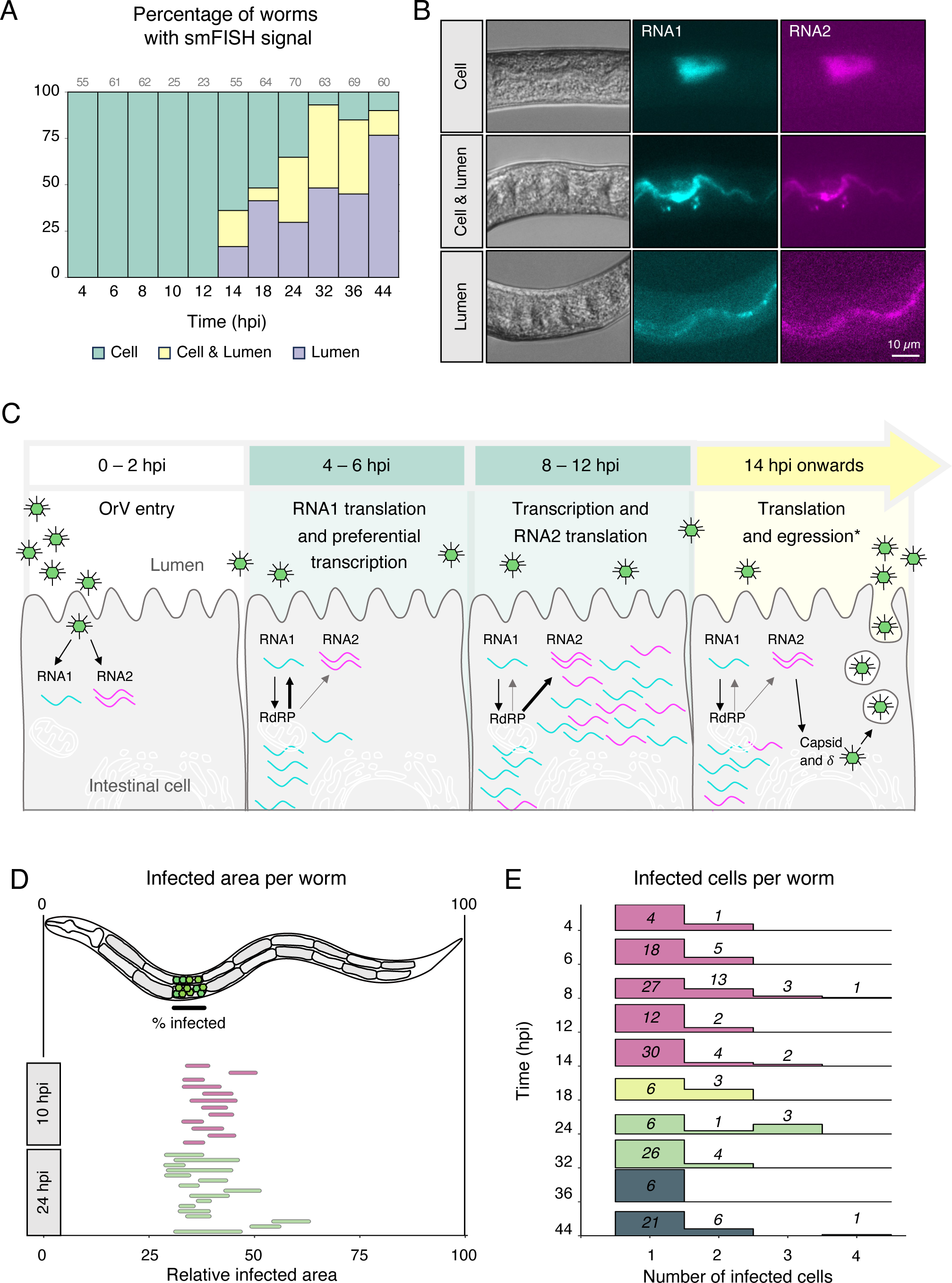
Spatial progression of the infection (A) Percentage of worms with OrV signal in intestinal cells, intestinal cells and luminal space or luminal space during development as determined by smiFISH. (B) Representative smiFISH images of worms with OrV in intestinal cells, intestinal cells and luminal space or luminal space at 24 hpi. (C) Schematic representation of our understanding of the OrV cycle. (D) Infected area per worm relative to the worm length as determined by smiFISH at 10 (pink; *n* = 12 worms) and 24 (green; *n* = 16 worms). (E) Number of infected cells per infected worm over time as determined by smiFISH. Density plot represents the relative abundance of the number of infected cells within each timepoint.

OrV has been described to colonize the anterior intestinal cells (Franz et al., 2014), but whether a late infection can also colonize posterior intestinal cells or whether the number of infected cells increases over time is still unknown. To answer these questions, we quantified the relative infected area per worm at 10 and 24 hpi (Fig. 2D). We did not observe a significant difference in the relative infected area at 10 *vs* 24 hpi (Mann-Whitney test, *P* = 0.3713), suggesting that the infection does not spread within the worm from the first viral load peak to the second viral load peak. We also quantified the total number of infected cells per infected worm during time and did not observe a significant difference at any time point (Fig. 2E; Kruskal-Wallis test, *P* = 0.3239), indicating that in most worms, throughout the viral replication cycle, OrV only colonizes, on median, 1 - 2 intestinal cell. Altogether, these results show that posterior colonization of the intestine does not take place within the observed timespan.

### *C. elegans* dynamic transcriptional response to OrV infection

To better understand the complexity of the host’s responses to a viral infection, we next focused on the transcriptomic response of *C. elegans* to OrV through time. For this purpose, we compared the transcriptome of control and infected worms at multiple time points until 44 hpi (Fig. 1A, Table S1). We performed a differential expression analysis (DEA) by time point which revealed a complex landscape (Fig. 3). We found a significant correlation (*rs* = 0.9000, 9 d.f., *P* = 0.0002) between the number of differentially expressed genes (DEGs) and OrV accumulation at each time point (Fig. 3A) with a predominance of upregulated genes in the first 18 hpi, and a more symmetric number of up- and down-regulated responses from 22 hpi on, although the strongest changes still correspond to overexpressed genes (Fig. 3A-C). Moreover, we identified a group of 29 upregulated DEGs at all timepoints with high viral loads (*i.e*., ≥ 0.5% of the total sample reads) (Fig. 3B, Fig. S3). All these DEGs were upregulated between 6 to at least 32 hpi; *col-133* was upregulated at every time point analyzed (Fig. 3C, Fig. S3B). Twenty-one of these 29 genes, including *eol-1*, *ddn-1* and the *pals* genes, were already characterized as part of the IPR (Reddy et al., 2019). Among the rest, *math-39* and *Y105C5A.9* were identified as dependent of the ZIP-1 transcription factor highly associated with IPR (Lažetić et al., 2022) and positive modulator of non-sense mediated mRNA decay (Kim et al., 2020). This is the first time that *col-133*, *trpl-5*, *math-15*, *Y43F8B.12*, and *Y6G8.5* have been associated to the worm’s response to OrV. A smaller group, formed by *col-133*, *ddn-1*, *eol-1*, *fbxa-182*, *math-15, pals- 33*, and *pals-37*, was differentially expressed from 6 hpi until the end of the time course (Fig. 3B, 3C). We also performed a functional enrichment analysis (FEA). The FEA was carried out on the up-and down-regulated genes separately and, contrary to what happened with DEGs, the number of significant categories in the downregulated responses was bigger than in the upregulated ones (Fig. 3D). Specifically, we found a first strong response between 12 and 18 hpi in the three Gene Ontologies (GO: Biological Process (BP), Molecular Function (MF) and Cellular Component (CC)) and in the metabolic data from KEGG and Wikipathways, and a second massive response around 26 hpi in all the databases explored (Fig. 3D, complete results in Table S2).

**Figure 3.**
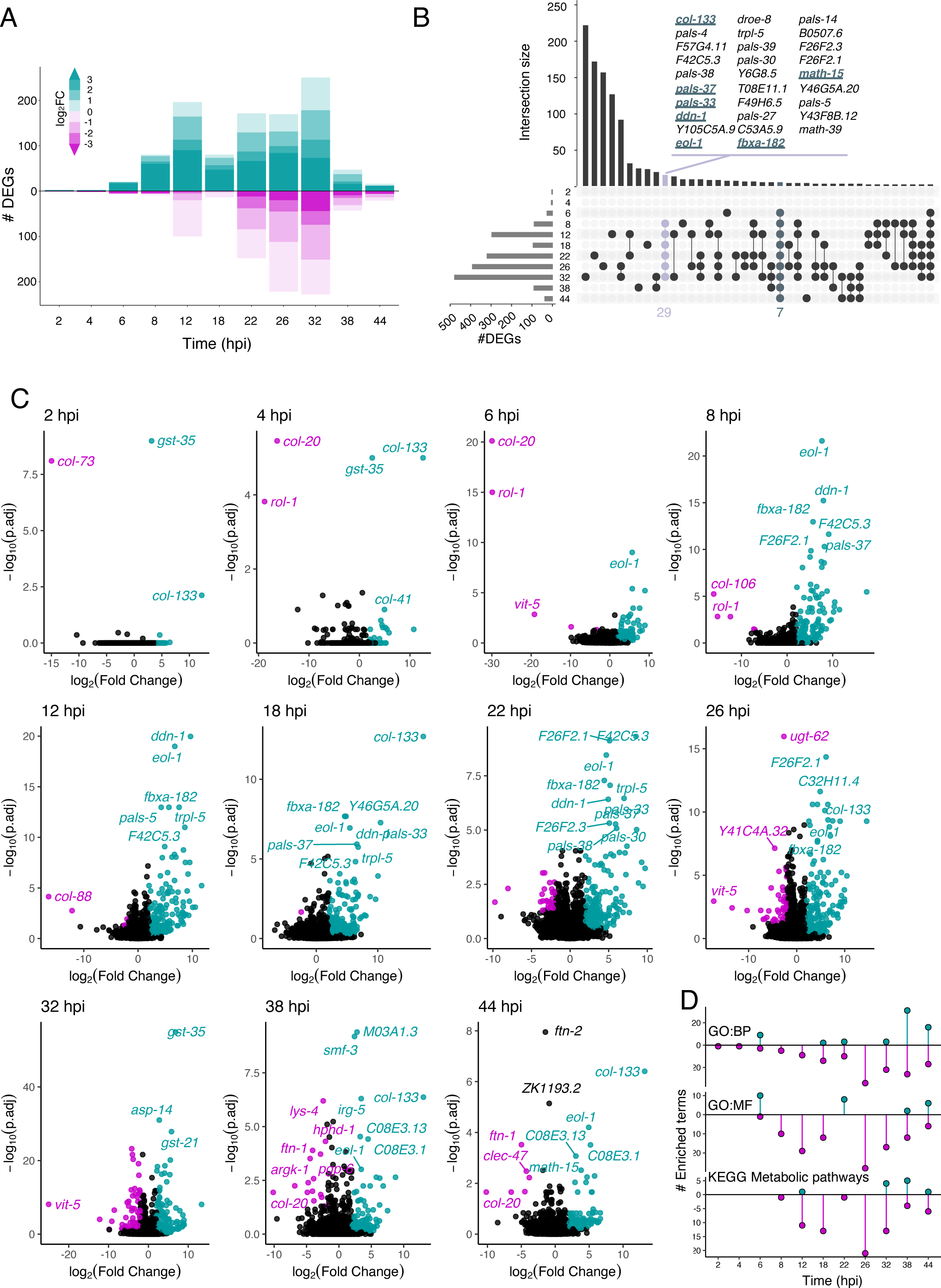
*C. elegans* transcriptomic responses across the OrV infection process (A) Number of significant DEGs (adjusted *P* ≤ 0.0499) up- (cyan) and down- (magenta) regulated at each point analyzed. The different intensities indicate the log2-fold change value. (B) Upset plot showing the intersections between the DEGS at each time. The points indicate the sets that are counted in the vertical bar. The horizontal bars indicate the total number of DEGs at each time. We found 29 genes ubiquitous to the exponential viral replication and host-pathogen conflict stages (*B0507.6*, *C53A5.9*, *<u>col-133</u>*, *<u>ddn-1</u>*, *droe-8*, *<u>eol-1</u>*, *F26F2.1*, *F26F2.3*, *F42C5.3*, *F49H6.5, F57G4.11*, *<u>fbxa-182</u>*, *<u>math-15</u>*, *math-39, pals-4*, *pals-5, pals-14*, *pals-27, pals-30*, *<u>pals-33</u>*, *<u>pals-37</u>*, *pals-38*, *pals- 39*, *T08E11.1*, *trpl-5*, *Y105C5A.9*, *Y6G8.5, Y46G5A.20*, and *Y43F8B.12*). Seven of them (underlined) persistent from 6 hpi to the end of the infection process. (C) Differentially expressed genes at each time point. Cyan and magenta represent significantly upregulated and downregulated genes, respectively. (D) Number of significant enriched terms (adjusted *P* ≤ 0.05) per time. The FEA was performed on the up and down DEGs separately. Gene Ontologies biological process (GO:BP) and molecular function (GO:MF) and KEGG metabolic pathways for *C. elegans* are shown. See also Fig. S2 and Fig. S3.

When focusing on the upregulated responses in the early stages of infection, at 6 hpi the following GO:BPs were enriched: nervous system development, axon extension involved in axon guidance, nuclear RNA surveillance and regulation of transcription, DNA-templated (Fig. S2A); and the GO:MFs related with 5’,3’-exonuclease activity and RNAII polymerase transcription factor activities (Fig. S2B). The TGF-β signaling pathway was upregulated at 12 hpi (Fig. S2C). Some of the responsible genes for this upregulation, *cul-6*, *skr-4* and *skr-5*, are skp1-cullin1-F-box (SCF) ubiquitin ligase components and part of the IPR (Reddy et al., 2017). Other genes involved in the TGF-β signaling pathway were identified as DEGs at different timepoints (*lin-35*, *mpk-1*, *rho-1*, *skr-3*, and *skr-19*) confirming the importance of this pathway throughout the infection.

When focusing on the early downregulated genes, we observed an earlier enrichment in significant terms. Within the first 2 hpi, there was a downregulation of genes involved in extracellular matrix organization; within 6 hpi in genes involved in collagen and cuticle development (Fig. S2A) and lipase activity (Fig. S2B); and within 8 hpi in genes related to helicase activities and UDP-glycosyltransferase activities (Fig. S2B). When analyzing the DEGs using KEGG pathways, we noticed an enrichment in other types of O-glycan biosynthesis at 8 hpi (Fig. S2C), a downregulation in mitochondrial translation and proteolysis at 12 hpi (Fig. S2A) and significant effects upon peroxisome, lysosome, metabolism of sphingolipids, pentose phosphate pathway, and ABC transporters (Fig. S2C).

During the host-pathogen conflict phase we observed upregulated genes related with the defense response to bacterium, the regulation of tube size and epithelial cell differentiation (Fig. S2A), as well as poly(A)- and poly(U)-RNA binding and receptor tyrosine kinase binding (Fig. S2B). In contrast, the downregulated genes were associated with the control of proteolysis by protein deneddylation (removal of ubiquitin-like protein), regulation of transport and stress response to Cd^++^, as well as fatty acid elongation (Fig. S2A). Towards the end of this phase, genes related to autophagy were also downregulated.

During the persistent residual infection phase there was an increase of significant terms among the overexpressed genes, including genes related with the regulation of MAPK cascade, Rho protein signaling transduction, mitochondrial unfolded protein response (UPR^mt^) and acetyl-CoA metabolic processes (Fig. S2A). Genes related with the glutathione metabolism were enriched from 38 hpi (Fig. S2C) and some categories previously identified as downregulated were upregulated in this phase, including fatty acid elongation and very long fatty acid metabolic process (Fig. S2A), pentose phosphate pathway and glycolysis/gluconeogenesis (Fig. S2C).

### DEGs that correlate or anticorrelate with the virus accumulation

This work provides, to our knowledge, the best transcriptomic temporal resolution of an OrV infection process. To reach more sensitivity, we performed a likelihood ratio test (LRT), identifying the genes with an expression profile altered by the infection (interaction between condition and time). This contrast revealed a total of 2313 DEGs (adjusted *P*-value ≤ 0.0499, Table S3), 1382 of which were not detected in the DEA shown in the previous section (Fig. 4A). For exploratory reasons, the genes were clustered by their profiles on 32 different groups (Fig. S4). When focusing in the expression pattern of the 150 most significant genes obtained using this contrast (Fig. 4B) we noticed they could be grouped into five different profiles (Fig. 4C - 4G), distinguishing between early upregulation (Fig. 4C), late upregulation (Fig. 4D, 4E) and late downregulation (Fig. 4F, 4G). Some of the genes highlighted above (*ddn-1*, *eol-1* and *ZC196.3*) and multiple *pals* (*pals-1*, -*3*, -*4*, -*5*, -*27*, -*30*, -*32*, -*37*, -*38*, and -*39*) share the early upregulation profile (Fig. 4C).

**Figure 4.**
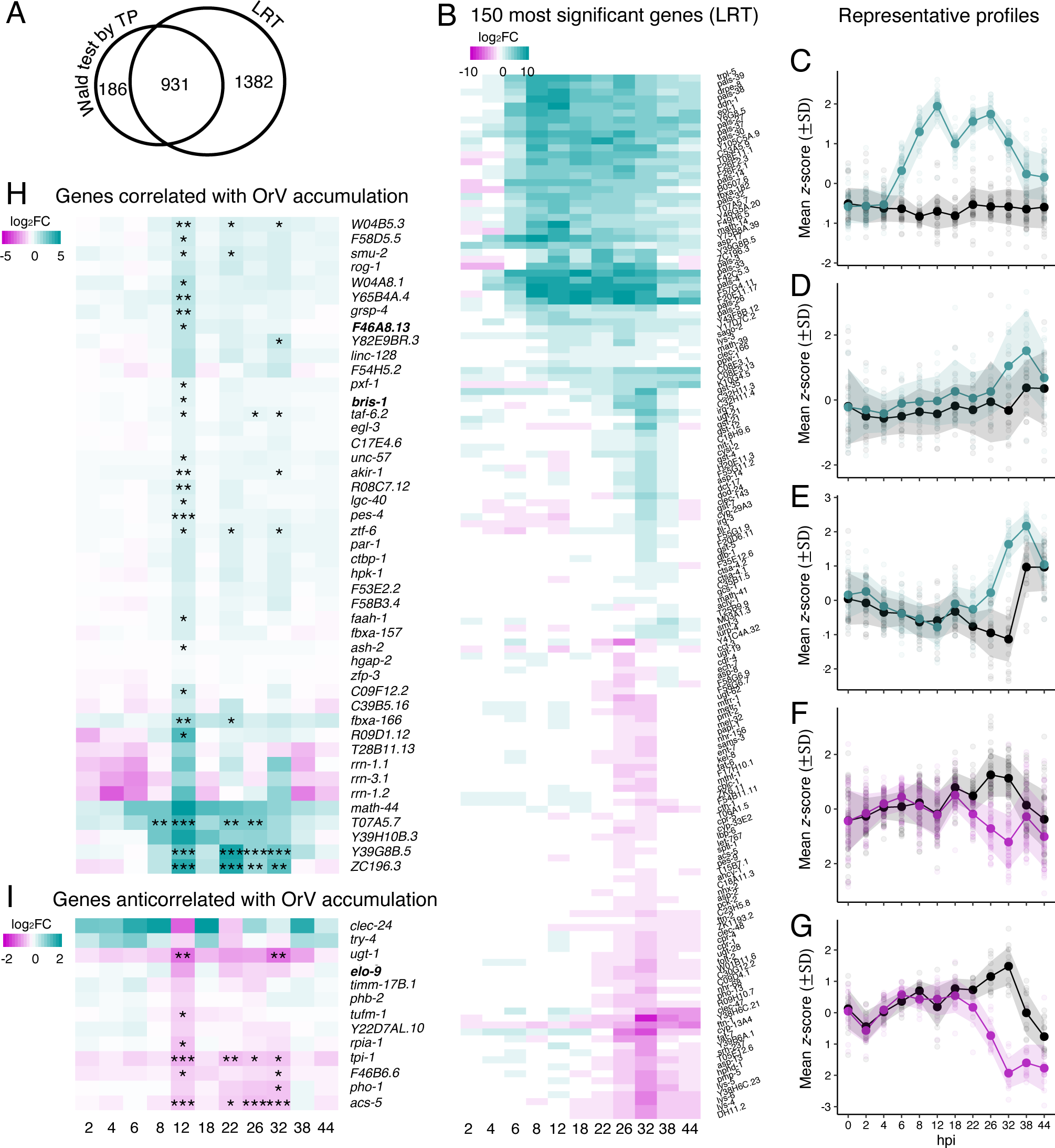
Differential gene expression (A) Total number of DEGs obtained by DEA individual time point (TP) Wald test and likelihood ratio test (LRT) to obtain those with an expression profile affected by the infection process. (B) Expression profile of the 150 most significant genes detected with the LRT. (C-G) Representative profiles of the 150 most significant genes. The mean *z*-score (±1 SD) of each group is represented by the line and the shadow respectively. Lighter points represent *z*-scores of the DESeq2 transformed values of each gene. Black represents the control and cyan or magenta represent the infected group and general up- or down-regulation of the cluster, respectively. (H) Expression profile of DEGS correlated (*r* ≥ 0.8011) with viral accumulation. In bold, genes with *r* ≥ 0.9148. (I) Expression profile of DEGs anticorrelated (*r* ≤ −0.8012) with viral accumulation. In bold, *r* = −0.9131. Asterisks indicate adjusted significance level at each time point: \**P* ≤ 0.05, \*\**P* ≤ 0.01 and \*\*\**P* ≤ 0.001. See also Fig. S4.

Furthermore, we identified the genes that correlate with OrV accumulation dynamics among all the differentially expressed ones. A total of 45 genes showed a highly positive correlation (*r* ≥ 0.8011, 9 d.f., *P* ≤ 0.0030) with OrV accumulation (Fig. 4H) while 13 presented an anticorrelated profile (*r* ≤ −0.8012, 9 d.f., *P* ≤ 0.0030) (Fig. 4I). Among the positively correlated genes, two had a (*r* ≥ 0.9148, 9 d.f., *P* < 0.0001): *bris-1*, enriched in neurons and predicted to enable guanyl-nucleotide exchange factor activity and to be involved in regulation of ARF protein signal transduction, and *F46A8.13*, a long non-coding RNA also enriched in neurons. In contrast, *elo-9*, a protein with fatty acid elongase activity, had a *r* = −0.9131 (9 d.f., *P* ≤ 0.0001). Remarkably, some of the positively correlated genes were already associated with OrV response: *Y39G8B.5* and *ZC196.3* are catalogued as part of the IPR (Reddy et al., 2019) and *C17E4.6*, *pes-4*, *pxf-1*, *taf-6.2*, *T07A5.7*, and *Y39G8B.5* are ZIP-1 dependent (Lažetić et al., 2022). Moreover, *clec-24*, a C-type lectin predicted to be associated with the membrane and an antimicrobial effector, displays a negative correlation with virus accumulation. Among the aforementioned, *T07A5.7*, *Y39G8B.5* and *ZC196.3* show the expression profile in Fig. 4C, while *acs-5* is a characteristic representant of the expression profile shown in Fig. 4G.

### Some of the identified responses are shared with other infections

After characterizing the worm’s responses to OrV infection on a temporal scale, we wanted to tease apart those specific to OrV infection from those common to other biotic stresses. For this purpose, we compared the DEGs in wild-type N2 worms upon infection with OrV, Gram− and Gram+ bacteria and fungi (Yang et al., 2016; Chen et al., 2017; Osman et al., 2018). A total of 55 experiments were included in the analysis (Fig. S5A, Table S4). We found 221 genes unique to the OrV transcriptomic response (OrV-specific genes from now) and 135 genes affected by all pathogens (Fig. 5A).

**Figure 5.**
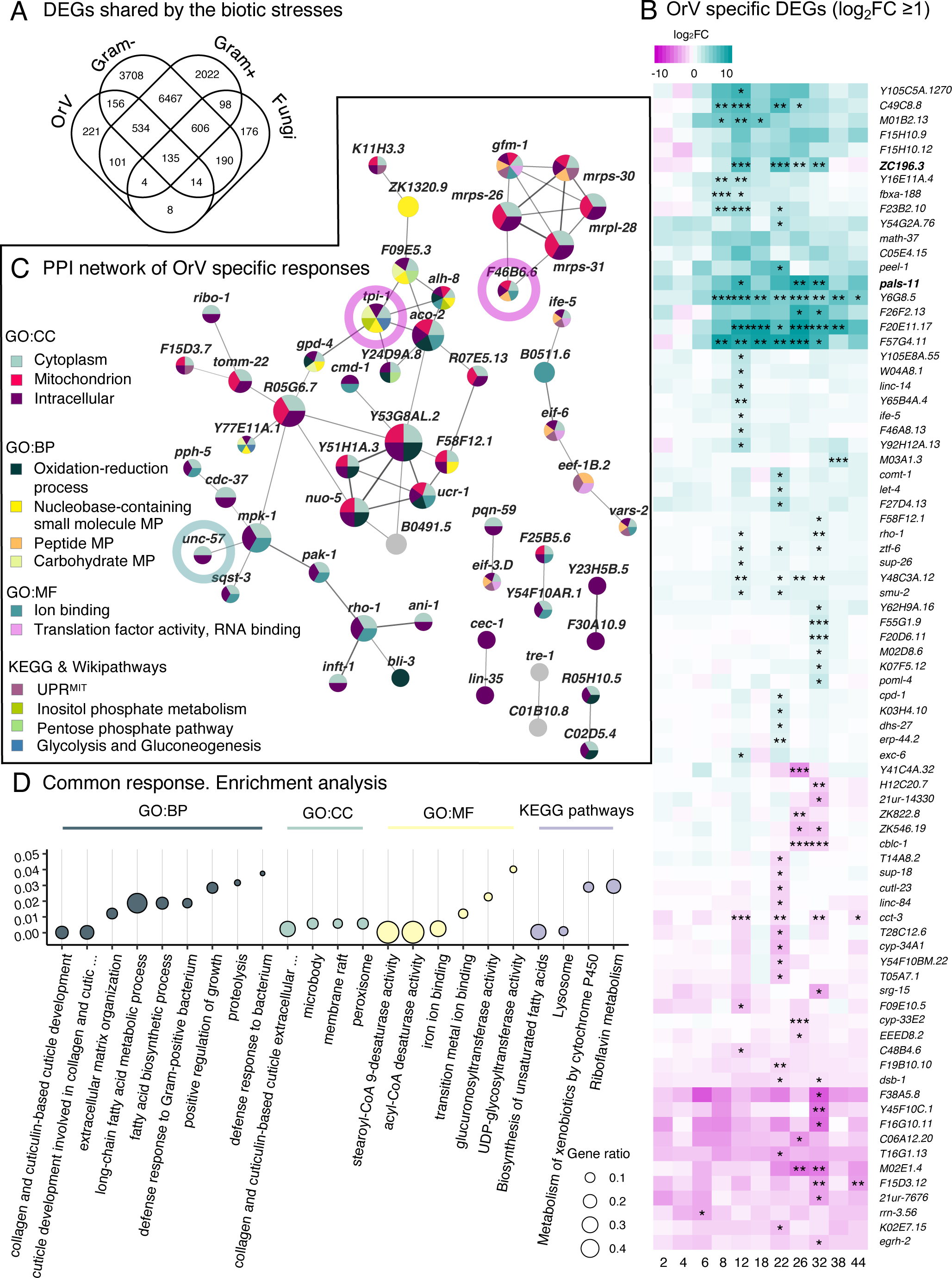
Comparison with other biotic stresses (A) Intersections between the different categories of stresses. A gene was included in a biotic stress category when was detected as DEG in any experiment belonging to the category. There are a group of 221 genes in the OrV response that do not appear in the other experiments included (only experiments in N2 were compared). 135 genes are common to all the categories studied. (B) Expression profile of OrV specific DEGs with a |log2-fold change| > 1 for at least one time point. Asterisks indicate adjusted significance level at each time point: \**P* ≤ 0.05, \*\**P* ≤ 0.01 and \*\*\**P* ≤ 0.001. (C) Protein-protein interaction network of the genes specific to OrV. Node color indicates enriched category (adjusted *P* < 0.05) and size indicates degree. Outer node circles indicate correlation (cyan) or anticorrelation (magenta) with virus accumulation. (D) Functional enrichment analysis of the genes common to all the stresses. See also Figs. S5-S8.

Within the OrV-specific responding genes, we noticed a variety of gene expression profiles (Fig. 5B). Some of the OrV-specific genes were already identified above: *ash-2*, *akir-1*, *F46A8.13*, *pxf-1*, *smu-2*, *taf-6.2, unc-57*, *W04A8.1*, *Y65B4A.4*, *ztf-6,* and *ZC196.3* are among those positively correlated to virus accumulation (Fig. 4H), while *F46B6.6* and *tpi-1* are anticorrelated to viral load (Fig. 4H); *F57G4.11* and *Y6G8.5* (Fig. 5B) are ubiquitous DEGs (Fig. 3B); and *F57G4.11*, *pals-11* and *ZC196.3* are classified as part of the ZIP-1-independent IPR (Lažetić et al., 2022). A functional enrichment of the OrV-specific genes highlighted the intracellular compartment, the cytoplasm and the mitochondria; and processes such as oxidation-reduction. The metabolic pathways inositol phosphate and the mitochondrial unfolded protein response (UPR^mt^) were also significantly enriched. Some of the categories enriched are shown on the protein-protein interaction network (PPIN) formed by the OrV- specific genes, including the modules formed by the mitochondrial proteins, which contains the anticorrelated gene *F46B6.6*, or the branch of the central module with the virus-correlated gene *unc-57* (Fig. 5C). Although genes traditionally highlighted as virus-affected are among those upregulated, the temporal expression pattern of the OrV-specific genes is quite variated (Fig. 5B), and those forming part of the PPIN shows a predominant downregulation (Fig. S5B).

Next, we asked whether these specific responses to viral infections were shared with other species of *Caenorhabditis*. To this end, we compared the DEGs upon infection of *Caenorhabditis briggsae* with Santeuil virus (Chen et al., 2017) with our results. We found that out of the 221 OrV-specific genes, 181 genes had homologs in *C. briggsae* and only *C49C8.8*, *F15H10.9*, *F23B2.10*, and *ZC196.3* were differentially expressed upon infection with both viruses (Table S5). *ZC196.3* has been described as part of the IPR, *F23B2.10* is affected by ZIP-1 (Lažetić et al., 2022), and *F15H10.9* is altered by infection with *Nematocida parisii* in *drh-1(ok3495)* (Reddy et al., 2019). *C49C8.8*, which is overexpressed at 8 hpi and has an expression pattern resembling the damping waves profile (Fig. S4, cluster 8), appears to be the most unique to viral responses.

Within the set of genes affected by all pathogens (Fig. 5A) we found some of our ubiquitous genes (Fig. 2B): *B0507.6*, *ddn-1*, *droe-8*, *F42C5.3*, *pals-27*, *pals-39*, and *Y46G5A.20*. Interestingly, they are also classified as part of the early ZIP-1-dependent IPR genes (Lažetić et al., 2022). The antimicrobial effectors *clec-51*, *clec-53*, *clec-67*, *clec-70, clec-72*, *clec-227*, *cnc-4*, *lys-2*, *lys-5*, and *lys-6*, are also among the common responses.

Next, we focused on three specific immune-related categories: IPR response, antimicrobial effectors and lipid metabolism. We found 65 out of the 80 IPR genes within our DEGs (adjusted *P* ≤ 0.0468) (Fig. S6A). Some of the IPR genes were also involved in the response to extracellular pathogens: 52.5 % of the IPR genes were differentially expressed 12 hpi with *Bacillus thurigiensis* and 50% at 24 hpi with *Enterococcus faecalis* (Yang et al., 2016) (Fig. S6B), suggesting that the role of IPR against diverse pathogens remains to be clarified.

We also noticed more antimicrobial effectors involved in antiviral responses than previously reported (Dierking et al., 2016): several C-type lectins were up- and down-regulated (Pees et al., 2021) (Fig. S7A), the lysozyme *lys-3* was upregulated at 18 and 26 hpi, *lys-2* at 32 hpi and *lys-4*, *lys-5* and *lys-6* were downregulated from 26 hpi. Moreover, the antimicrobial effectors *clec-51, clec-53, clec-67, clec-70, clec-72, clec-227, cnc-4*, *lys-2,* and *lys-6* were differentially expressed upon OrV, Gram+ and Gram– bacteria, and fungi infections (Fig. S7B).

The lipid metabolism plays key roles in a variety of diseases, whether pathogenic in origin or not (Yoon et al., 2021). The feedback between of OrV replication and lipid metabolism has been already studied (Casorla-Perez et al., 2022b). Here we found almost 35% of the genes associated with lipid metabolism (GO0006629) in our DEGs list (Fig. S8A). Of these, *hyl-1*, *pcyt-2.1*, *psd-1*, *scp-1*, and *T26C12.1* were amongst the OrV-specific genes and *hyl-1* and *scp-1*, which were significantly upregulated at 18 and 22 hpi respectively, are predicted to be regulatory genes. A general trend towards downregulation was observed, as previously described (Casorla-Perez et al., 2022a). However, the biosynthesis of unsaturated fatty acids was one of the most significant categories enriched in the common genes (Fig. 5D). Of note, the metabolism of lipids is also important in the responses to other biotic stresses (Fig. S8B) and *dld-1*, *fat-2*, *fat-5*, *fat-6,* and *sptl-2* are part of the common response to all pathogens tested so far.

### Pro- and anti-viral functions of DEGs

We finally sought to characterize the role of some of the DEGs upon OrV infection. We selected the secondary argonautes *ppw-1*, *sago-2* and *vsra-1*, which were differentially upregulated early in infection (Fig. S9A). We also selected *lys-2*, a lysozyme involved in the defense response to bacteria and fungi, and the antimicrobial peptide *clec-60*, due to their roles in the defense response against microorganisms (Fig. S9A). Additionally, we selected *gst-35*, a glutathione transferase which may have roles in detoxification and cell signaling; *eol-1*, an RNA pyrophosphohydrolase involved in the regulation of olfactory learning; and *C53A5.9*, a paralog of influenza virus NS1a binding protein homolog, because they were significantly upregulated during early infection (2 to 6 hpi) (Fig. S9A). Finally, we included *drh-1* and *pals-22* mutants as controls for sensitivity and resistance, respectively (Ashe et al., 2013; Reddy et al., 2019).

To assess the role of these proteins upon OrV infection we studied whether the absence of these proteins would impact viral accumulation and infectivity. *vsra-1(tm1637)*, *clec-60(tm2319)*, *ppw- 1(pk2505)*, and *sago-2(tm894)* were ordered from the CGC whilst *C53A5.9(tm10539)*, *eol-1(tm6609)* and *lys-2(tm2398)* were ordered from the NBRP and backcrossed three times to a wild-type N2 background. *drh-1(ok3495)* was a gift from M.A. Félix and *pals-22(jy81)* from E.R. Troemel. A *gst-35* mutant, *gst-35(esv1)*, was generated using CRISPR/Cas9 and resulted in a 478 bp deletion which included part of the first exon and a splice junction, resulting in a truncated protein which only maintained homology to the first three amino acids (Fig. S9B-E).

We first examined the impact of these proteins on OrV infectivity by looking at three key factors: the percentage of infected worms (Fig. 6A and 6B), the RNA2 signal positive worms (Fig. 6C), and the number of infected cells at 12 hpi (Fig. 6D), which aligns with the initial peak of viral load in wild-type worms (Fig. 1B). For infectivity, the fit of the data to a logistic regression revealed significant differences among genotypes (χ^2^ = 394.9264, 10 d.f., *P* < 0.0001). Further Bonferroni sequential *post hoc* test identified five distinct groups (adjusted *P* ≤ 0.0395). Mutants *clec-60*, *gst-35*, *lys-2*, and *sago- 2* showed no significant difference from the wild-type. By contrast, *pals-22(jy81)* and *ppw-1(pk2505)* formed a group with an ∼80% reduction in infectivity, while *eol-1(tm6609)* and *vsra-1(tm1637)* had a significant but smaller reduction (33.92% and 50.47%, respectively). Conversely, *drh-1(ok3495)* was 17.96% more susceptible to infection.

**Figure 6.**
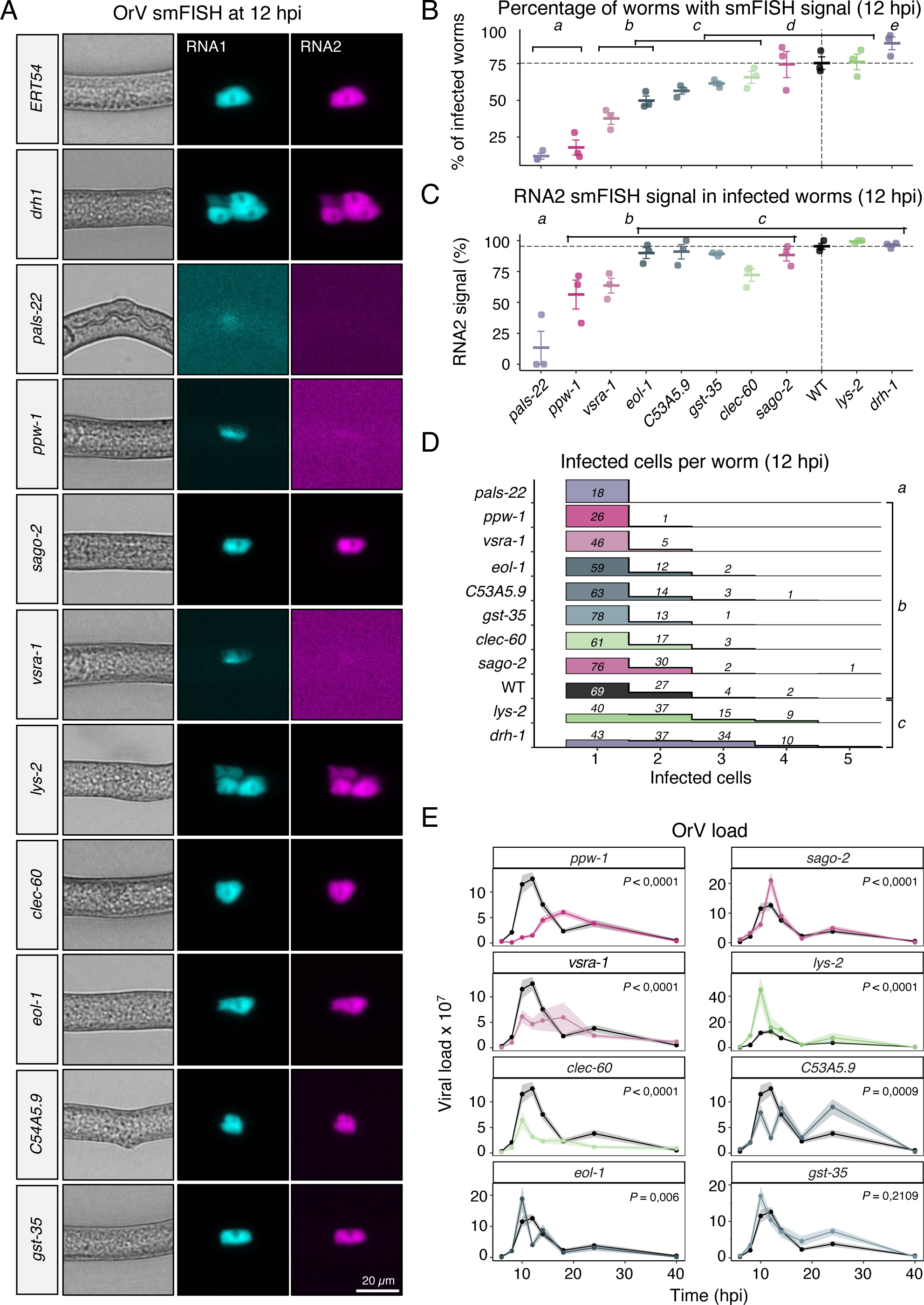
OrV dynamics in mutants of identified DEG (A) Representative smiFISH images of WT, *pals-22*, *ppw-1*, *vsra-1, eol-1*, *C53A5.9*, *gst-35*, *clec-60*, *sago-2*, *lys-2* and *drh-*1mutants at 12 hpi. (B) Percentage of infected worms at 12 hpi as determined by smiFISH. *n* = 49 ±5 per point. Error bars represent SD. (C) Ratio of RNA2/RNA1 positive worms determined by smiFISH in the indicated mutants. *n* = 27 ± 12 per point. Error bars represent SD. (D) Number of infected cells per infected worm over time as determined by smiFISH. Density plot represents the relative abundance of the number of infected cells within each mutant. (E) Viral load (±1 SEM; *n* = 5 replicates per time point) determined by RT-qPCR at 6, 8, 10, 12, 14, 18, 24, and 44 hpi in *ppw-1*, *sago-2, vsra-1, lys-2*, *clec-60, C53A5.9*, *eol-1* and *gst-35* mutants (colored lines and dots) and WT (black lines and dots). *P* values correspond to the result of the sequential Bonferroni *post hoc* pairwise test comparing the temporal dynamics of OrV accumulation on each mutant with the wild-type worms. See also Fig. S9.

Regarding the RNA2 positive worms, significant differences were found among genotypes (Fig. 6C, logistic regression: χ^2^ = 176.7448, 10 d.f., *P* < 0.0001). Three significant groups emerged (*post hoc* adjusted ≤ *P* 0.0473). *pals-22(jy81)* had an 88.36% lower accumulation of RNA2 compared to wild-type relative to RNA1. *ppw-1(pk2505)* and *vsra-1(tm1637)* showed a moderate reduction (∼36%). The rest of the mutants showed no significant difference from wild-type.

We then analyzed the distribution of infected cells per worm (Fig. 6D). Due to non-normal distribution and heteroscedasticity of the data, a Kruskal-Wallis test was used, showing significant differences among genotypes (χ^2^ = 184.4403, 10 d.f., *P* < 0.0001). Games-Howell *post hoc* analysis revealed three distinct groups (adjusted *P* ≤ 0.0473). At the one side, *pals-22(jy81)* had only one infected cell per worm, significantly less than the wild-type. At the other side, *lys-2(tm2398)* and *drh- 1(ok3495)* had roughly 53% more infected cells than wild-type worms.

Together, these results suggest LYS-2 and DRH-1 prevent multicellular OrV colonization and PALS-22, PPW-1 and VSRA-1, which had lower infectivity, a reduction in RNA2 expression relative to RNA1, and fewer infected cells per worm, hinder OrV amplification early in viral replication. The other mutants behaved like the wild-type in these analyses.

To enhance our comprehension of these proteins’ roles in OrV infection, we proceeded to examine viral accumulation in the mutants. We monitored the progression of viral load in synchronized mutant populations from hatching to the end of larval development (L4 stage). OrV inoculation took place upon hatching, and samples were collected at various hpi for RT-qPCR analysis (Fig. 6E). The data from the accumulation curves were fitted to a generalized linear model with a gamma distribution and log-link function. Worm genotype and hpi were considered as model factors. We specifically looked for differences in the temporal dynamics of OrV accumulation between mutant and wild-type worms and conducted sequential Bonferroni *post hoc* pairwise tests. We observed significantly higher OrV accumulation in *lys-2(tm2398)* and *sago-2(tm894)* (*P* < 0.0001), with *lys-2(tm2398)* showing a 4-fold increase in viral load at 10 hpi, and *sago-2(tm894)* exhibiting a 1.6-fold increase at 12 hpi. This suggests that these genes may play a direct role in the antiviral response. Conversely, we noted significantly lower OrV accumulation in the other secondary argonautes, *ppw-1(pk2505)* and *vsra-1(tm1637)* (*P* ≤ 0.0001), with viral load peaks at 18 hpi. This trend was also observed in the antimicrobial peptide *clec- 60(tm2319)* (*P* < 0.0001), along with a disrupted OrV accumulation pattern in *C53A5.9(tm10539)* and *eol-1(tm6609)* (*P* ≤ 0.0009). Lastly, we did not detect differential OrV accumulation in *gst-35(esv1)* (*P*= 0.2109), suggesting that its altered expression upon infection may be an indirect effect unrelated to the defense response. However, we cannot rule out the possibility that redundancy within the glutathione transferase family might be masking a potential role in OrV infection.

From these experiments we conclude that *C53A5.9*, *clec-60*, *eol-1*, *ppw-1*, and *vsra-1* had proviral functions, whilst *lys-2* and *sago-2* appear to have antiviral functions. *gst-35* appears to be either redundant or not involved in the defense response.

## DISCUSSION

### OrV dynamics along larval development

In this study we followed the progression of a viral infection within a population which was synchronously infected. Regardless the quantification method employed, we observed that the infection was highly dynamic and followed four phases: (1) a pre-replication phase in which the viral load and percentage of infected worms remained low and RNA1 was preferentially transcribed, which lasted ∼6 hours; (2) an exponential viral replication phase in which the viral load and the number of infected worms rapidly increased, spanning from 8 to 14 hpi; (3) a phase of host-pathogen conflict in which the percentage of infected worms and the viral load remained fairly constant, from 16 to 36 hpi; and (4) a phase of persistent residual infection from 38 to 44 hpi. The ratio of RNA2 to RNA1 was dynamic throughout the infection and started with ∼2 RNA2 molecules per RNA1 molecule, which opens the possibility that more RNA2 gets encapsulated in each virion. Moreover, RNA1 preferentially accumulated during the initial part of the exponential viral replication phase. During the second part of the exponential viral replication phase RNA2 levels increased up to four times more than RNA1 levels, suggesting that the structural proteins encoded in RNA2 were needed to complete the viral replication cycle and egression. In agreement with these results, we started observing OrV smiFISH signal in the lumen at 14 hpi, which allowed us to determine the beginning of egression at around 14 hpi. Altogether, our data suggests that OrV accumulation during larval development is highly dynamic and that a viral replication cycle would be complete at 14 hpi, which would be consistent with the increase in RNA2 levels leading up to 12 hpi and the decrease in viral load after 12 hpi.

Moreover, we observed that the number of infected cells per worm remained close to one throughout the rest of the infection, suggesting that the increase in viral load we observed by RT-qPCR and RNA-seq is due to intracellular viral replication, possibly in the same infected cells. The ability of neighboring cells to ward off the virus raises the possibility that the infected cells share some diffusible signal with their neighbors to induce an antiviral state, as has been observed in mammalian cells (Song et al., 2023). In *lys-2* and *drh-1* mutants the number of infected cells was increased, as was the viral load, suggesting that the increase in viral load may be due to an increased number of cells and that LYS- 2 and DRH-1 prevent multicellular colonization. Another member of the RNAi pathway, RDE-1, is also necessary to prevent multicellular colonization (Guo et al., 2017), hinting that DRH-1 may prevent multicellular colonization due to its role in the RNAi pathway and not through activating the IPR response. Lastly, the viral load in *lys-2(tm2398)* was reduced to wild-type levels after the initial peak, suggesting that *lys-2* is involved in preventing multicellular colonization but unlike *drh-1*, is not involved in intracellular defense.

We also observed that the relative infected area per worm did not increase over time, which may be due to the differential localization of anti-viral genes within the intestine. It has been recently reported that intestinal cross-linking collagens, which are restricted to the posterior intestinal cells, mediate antiviral immunity in the intestine (Zhou et al., 2023), with intestinal RNAi of different collagens resulting in increased viral loads. This increase in viral load may thus be due to an increased number of infected cells, as we observed in the *lys-2* and *drh-1* mutants.

By analyzing the transcriptomic response during the whole span of larval development we have been able to characterize temporal responses to OrV infection. We identified a group of genes upregulated through the whole infection and genes whose expression profiles were correlated or anticorrelated with viral accumulation. A comparative analysis of the responses elicited by other biotic pathogens in wild-type worms teased apart OrV-specific genes from defense genes responding in a non-pathogen specific manner.

Finally, the developmental stage of the host has an impact on the response to viral infections (Melero et al., 2023). Our experiments were performed in L1 larvae, unlike a previous study in which the transcriptomic response to OrV in wild-type worms was analyzed in L3 larvae (Chen et al., 2017). At 12 hpi we see some key differences, such as the absence of molting-related genes in the L3-infected dataset. This is probably because in their case the L3 molt had already passed, and in our case the 12 hpi was close to the L1 molt. Taking into consideration the developmental timing of the worms in future experiments will help us understand the effect of the viral infection in development and distinguish specific responses to OrV.

### The nervous system responds to OrV at the earliest stages of the infection

The communication between the nervous system and the intestine during an infection is essential for a proper immune response (Zhang and Zhang, 2009; Martineau et al., 2021; Wibisono and Sun, 2021; Liu et al., 2023), avoidance of infected food sources (Liu et al., 2020; Pees et al., 2021) and preferential mating behavior of the males with non-infected hermaphrodites (van Sluijs et al., 2021). Here, genes related with the nervous system, such as *eol-1*, play an important role from the first stages of the infection: absence of EOL-1, which is related to olfactory learning, has an impact on the viral load profile, with a higher maximum viral load than the control at an earlier time point. Moreover, a subsequent decline in viral load suggests an earlier activation of the defense response, as the infection process continues similarly than that of wild-type worms. The involvement of *eol-1* in the response to the virus is unclear but our results suggest that it contributes to shape the process, possibly due to erratic behavior that fails to avoid the virus.

### Influence of OrV on lipid metabolism, the UPR^mt^ and the inositol phosphate pathway

Our study unveiled a dynamic pattern of immune, stress, and metabolic responses throughout the OrV infection process. While extensive research on the impact of viral infections on metabolism has been conducted in mammals (Li et al., 2023), studies in *C. elegans* have predominantly focused on bacterial infections (Anderson and Pukkila-Worley, 2020).

Viral infections impact the lipid metabolism in several pathosystems: some viruses trigger the accumulation of lipid droplets while others reduce the lipid storage (Qu et al., 2023). OrV infection leads to a significant reduction in lipid accumulation, even though OrV itself relies on lipids for effective infection (Casorla-Perez et al., 2022). Our data revealed a general downregulation of lipid metabolism genes from 22 hpi, with downregulation of some responses as early as 12 hpi. Given that wild-type worms are capable of reducing the viral load, the reduction of the lipids could be modulated by the virus –as it is the case in other pathosystems (Heaton and Randall, 2011; Ketter and Randall, 2019; Herrera-Moro Huitron et al., 2023)– or could be part of an active response of the worm. Certainly, the fatty acid oleate, generated by *fat-6* and *fat-7*, plays a pivotal role in activating the innate system of *C. elegans* (Anderson et al., 2019). This significance is underscored by the inclusion of *fat-6* as one of the DEGs shared by biotic stresses in all the categories tested. This implies that *C. elegans* might have the capacity to adjust its lipid metabolism as a component of an engaged antiviral defense. Nevertheless, additional research is warranted to delve deeper into this phenomenon and gain a more comprehensive understanding.

Our study highlights the significant impact of viral infection on mitochondria, a crucial cellular component for maintaining balance through pathways like the unfolded protein response (UPR^mt^). This activation is vital for managing stress (Deng et al., 2019; Kumar et al., 2023). Under normal conditions, cytosolic ATFS-1 is transported into the mitochondria via the TOM complex for degradation. If this translocation fails, ATFS-1 enters the nucleus, triggering the expression of genes associated with mitochondrial chaperones, proteolysis, glycolysis, and innate immunity, such as *lys-2* (Martineau et al., 2021), which we found to have antiviral effects. Although we observed an enrichment of UPR^mt^ at 38 hpi, coinciding with the onset of the residual infection phase in which viral load decreased significantly, several indications point to an earlier pathway activation: (*i*) genes linked to mitochondrial translation were downregulated at 12 hpi; (*ii*) components of the TOM complex, *tomm-22* and *tomm-70*, were also downregulated at 12 and 26 hpi respectively, hindering ATFS-1 translocation; and (*iii*) ZIP-3, known to inhibit UPR^mt^ activation (Deng et al., 2019) displayed reduced expression (cluster 5 in Fig. S4). In contrast, ZIP-1, identified as a key regulator in the IPR (Lažetić et al., 2022), showed an upregulated expression profile (cluster 9 in Fig. S4). The expression patterns of both bZIP transcription family members align with the activation of immune genes through both pathways.

Among the enriched category of OrV-specific responses, we found the inositol phosphate metabolism. Within this category, specific DEGs like *alh-8*, *C54E4.5*, *sac-1*, and *tpi-1* connect inositol phosphate metabolism to various pathways including glycolysis/gluconeogenesis, tricarboxylic acid cycle, pentose and glucuronate interconversions, and glycerophospholipid pathways. These pathways were demonstrated to be enriched in our time-course experiment. Notably, *sac-1* is of particular interest, as it is predicted to facilitate the dephosphorylation of phosphatidylinositol-4-phosphate (PI4P), an antagonist of the PI4 kinase. The balance between this phosphatase and PI4K regulates PI4P concentration, influencing cell physiology, cell-signaling pathways, actin remodeling, membrane dynamics, and the life cycle of certain positive-stranded RNA viruses (Beziau et al., 2020). Specifically, a lipid environment rich in PI4P is crucial for the replication of enteroviral and flaviviral RNA synthesis (Hsu et al., 2010; Casorla-Perez et al., 2022b). SAC1 has been identified as a crucial host factor in regulating hepatitis B virus (Popescu et al., 2022; Zheng et al., 2023) and is pivotal for autophagosome-lysosome fusion (Zhang et al., 2020). This underscores *sac-1* as a promising candidate for further analysis.

### Proviral properties of *CLEC-60*

C-type lectins have been described to recognize and combat pathogens (Miltsch et al., 2014; Pees et al., 2021) and to also have pro-pathogenic functions. In fact, some RNA viruses, such as hepatitis A virus, influenza A viruses (Yang et al., 2023) and human immunodeficiency virus (Curtis et al., 1992; Geijtenbeek et al., 2000), bind to C-type lectins on the host cell surface to gain entry into target cells. Binding to C-type lectins is usually dependent upon the presence of glycosylated chains on the surface of the viral capsid. In *C. elegans*, C-type lectins are differentially regulated by bacterial and fungal infections and have functions related to feeding behavior, bacterial binding and antimicrobial capacities. Here we observed 47 out of 264 C-type lectins differentially expressed upon OrV infection. *clec-60*, which is expressed in the intestine, is differentially expressed upon bacterial of fungal exposure and RNAi knockdown leads to hyper-deformed anal region and severe constipation (O’Rourke et al., 2006; Irazoqui et al., 2010; Pees et al., 2015). Relevantly, *clec-60* is involved in antimicrobial defense against *Staphylococcus aureus* and *Microbacterium nematophilum* (O’Rourke et al., 2006; Irazoqui et al., 2010), among others. Our results indicate that *clec-60* is also involved in viral infection in *C. elegans*, and may have proviral roles, as *clec-60(tm2319)* had lower viral loads. These results suggest that similarly to what was observed in other RNA viruses, C-type lectins may be involved in OrV entry into host cells. However, further studies on the role of *clec-60* and other C-type lectins in OrV infection would be necessary to clarify this.

### Antiviral functions of *LYS-2*

Lysozymes form part of the innate immune response and the digestive system (Boehnisch et al., 2011; Gravato-Nobre et al., 2016). Lysozymes are mostly considered antimicrobial enzymes, but some reports have also described antiviral properties (Ferrari et al., 1959; Lee-Huang et al., 1999), including a recent study which found that lysozyme reduced the entry of SARS-CoV-2 in human epithelial cells (Song et al., 2022). In *C. elegans*, lysozymes have not been reported to be involved in the defense response against viruses (Dierking et al., 2016). Here we observed differential expression of 11 lysozymes upon OrV infection, indicating that lysozymes are regulated by viral infections in *C. elegans*. Moreover, some lysozymes were differentially expressed within hours of viral inoculation, including *lys-2*, which was upregulated within the first 2 hpi. Interestingly, *lys-2* was previously described to be regulated by all microbes it was exposed to except by OrV and *Yersinia pseudotuberculosis* (Dierking et al., 2016). Our results indicate that *lys-2* is also regulated by OrV. We did not observe a higher percentage of infected worms in *lys-2(tm2398)* when compared to wild-type worms, possibly due to the highly concentrated viral stock used in these experiments. However, we observed a significant increase in *lys-2(tm2398)* viral load upon OrV infection –with a viral peak four times higher than in wild-type worms– suggesting that *lys-2* has antiviral effects in *C. elegans*. Moreover, we observed a significant increase in the number of infected cells at 12 hpi, suggesting that *lys-2* prevents multicellular colonization. The decrease in viral load in *lys-2* mutants after the viral load peak suggests that the intracellular defense response which is able to clear OrV after colonization is not affected in *lys-2* mutants, suggesting that the anti-viral properties of *lys-2* rely in the prevention of multicellular colonization and not in intracellular defense mechanisms.

### OrV dynamics in argonaute mutants

Argonaute (AGO) proteins are highly specialized RNA-binding proteins that form complexes with small RNAs to direct RNA silencing. In *C. elegans* there are 27 AGO-like genes (Seroussi et al., 2023) and two of them, *rde-1* and *sago-2*, are necessary in the defense response against OrV: RDE-1 targets the OrV RNA and recruits the RdRP RRF-1 to generate secondary 22G-RNAs (Tabara et al., 1999; Félix et al., 2011; Sarkies et al., 2013) and SAGO-2 is thought be loaded with the secondary 22G-RNAs that were generated after targeting of OrV RNA by RDE-1 (Ashe et al., 2013; Seroussi et al., 2023) to target their degradation. In contrast, loss of AGO protein ALG-1 resulted in reduced OrV levels (Cubillas et al., 2023), suggesting that ALG-1 may have a role during the replication stage of OrV. Here we studied the role of the AGOs which were differentially expressed upon OrV infection (SAGO-2, PPW-1 and VSRA-1) all of which were overexpressed. Analysis of the AGO mutants upon OrV infection suggested that these AGOs have different functions during the infection. *sago-2(tm894)* showed a higher viral load coinciding with the 12 hpi viral load peak, confirming that SAGO-2 is necessary in the defense response against OrV. We did not observe a higher percentage of infected worms in *sago-2(tm894)* when compared to wild-type worms, possibly due to the saturating viral stock used in these experiments. In contrast, *ppw-1(pk2505)* and *vsra-1(tm1637)* showed reduced viral loads, percentage of infected worms and RNA2 to RNA1 ratios. These results suggest that PPW-1 and VSRA- 1 are necessary for viral replication and that viral replication in these mutants is stalled after the beginning of RNA1 replication but before RNA2 replication, similarly to what was observed with ALG- 1 (Cubillas et al., 2023), PALS-22 (Reddy et al., 2019) and *Nicotiana benthamiana* AGO1 (Pollari et al., 2020). However, it could also be that the loss of PPW-1 and VSRA-1 cause a misregulation of the immune system which benefits the host and results in reduced OrV levels. Indeed, PPW-1 regulates immune-responsive genes (Seroussi et al., 2023). Interestingly, even though the function of PPW-1 and SAGO-2 seems drastically different in regards to the defense response to OrV, both proteins have 98.65% amino acid similarity.

## Concluding remark

*C. elegans* stress responses often share common ground, and a single stressor can trigger various pathways. Consider, for instance, a proteotoxic stress that kickstarts not just the innate immunity, but also sparks a heat shock response and activates the UPR^mt^. Moreover, the developmental stage of the host can influence the nature of these responses. While it may sometimes seem like untangling a tightly woven knot, our work has shed a little light into the intricate sequences that dictate how the worm tackles an OrV infection at each stage of the process. This research offers a new lens through which to appreciate the nuanced strategies these tiny creatures employ in their battle against viral intrusion.

## ACKNOWLEDGMENTS

We thank Francisca de la Iglesia and Paula Agudo for excellent technical support and the members of the EvolSysVir lab for valuable comments and fruitful discussions. We thank M.A. Félix for strains JU2624 and RB2519 and for OrV JUv1580 and E.R. Troemel for strains ERT54 and ERT288. We also thank Wormbase (Davis et al., 2022). Some strains were provided by the Caenorhabditis Genetics Center, which is funded by NIH Office of Research Infrastructure Programs (P40 OD010440). Computations were performed on the HPC cluster Garnatxa at the Institute for Integrative Systems Biology (I2SysBio). This work was supported by grants PID2022-136912NB-I00 funded by MCIN/AEI/10.13039/501100011033 and by “ERDF a way of making Europe”, and CIPROM/2022/59 funded by Generalitat Valenciana to S.F.E. V.G.C. was supported by grant FJC2021-047264-I funded by MCIN/AEI/10.13039/501100011033 and by NextGenerationEU/PRTR. M.J.O-U. was supported by grant FPU2019/05246 funded by MCIN/AEI/10.13039/501100011033 and by “ESF investing in your future”. J.C.M-S. was supported by grant ACIF/2021/296 from Generalitat Valenciana. A.V-G. was supported by the European Space Agency contract 4000135960/21/NL/GLC/my to S.F.E.

## AUTHOR CONTRIBUTIONS

V.G.C., M.J.O-U, J.C.M-S, and S.F.E. conceived the study and designed the experiments. V.G.C. performed the experiments and analyzed data. M.J.O-U. performed the bioinformatic analysis and wrote the code. S.F.E. analyzed data. A.V-G. performed RNA extractions. V.G.C., M.J.O-U. and S.F.E. wrote the manuscript. All authors approved the latest version of the manuscript.

## STAR METHODS

### Resource Availability

#### Lead Contact

Further information and requests for resources and reagents should be directed to and will be fulfilled by the lead contact, Santiago F. Elena.

#### Materials availability

Materials are available from the lead contact upon request.

#### Data and code availability

RNA- seq data was deposited in the NCBI SRA: PRJNA

The necessary code to replicate the process can be found in https://git.csic.es/sfelenalab/OrV_progression.

Any additional information required to reanalyze the data reported in this work paper is available from the lead contact upon request.

#### *C. elegans* strain maintenance

All *C. elegans* strains used in this study are derived from the N2 Bristol strain unless specified, and are listed in the Key Resources Table. All strains were maintained at 20 °C on nematode growth medium (NGM) plates seeded with *Escherichia coli* OP50 bacteria under standard conditions (Brenner, 1974; Stiernagle, 2006). Mutants and transgenic strains were backcrossed three times into the N2 strain used in the laboratory unless already backcrossed into N2.

To achieve synchronized worm populations, plates containing eggs were meticulously rinsed with M9 buffer to eliminate larvae and adult worms while retaining the eggs. After a 1 h interval, plates were subjected to another M9 buffer wash to gather the larvae that had hatched during that period.

#### OrV stock preparation, quantification and inoculation

JU2624 or SFE2 worms coming from one freshly starved plate were inoculated with OrV by soaking in viral stock kindly provided by Prof. M.A. Félix (Félix et al., 2011) for 1 h. The volume was then divided among 30 9 cm NGM plates and allowed to grow for 5 d until starved. Worms were then resuspended in 15 mL of M9 (0.22 M KH2PO4, 0.42 M Na2HPO4, 0.85 M NaCl, 0.001 M MgSO4), let stand for 15 min at room temperature, vortexed and centrifuged for 2 min at 400 g. The supernatant was centrifuged twice at 21000 g for 5 min and then passed through a 0.2 μm filter.

RNA of the resulting viral stock was extracted using the Viral RNA Isolation kit (NYZ tech). The concentration of viral RNA was then determined by RT-qPCR (details below) and normalized using a standard curve. Primers used can be found in Table S6.

For the standard curve cDNA of OrV was obtained using Accuscript High Fidelity Reverse Transcriptase (Agilent) and reverse primers at the 3’ end of the virus. Approximately 1000 bp of the 3’ end of RNA1 and RNA2 were amplified using forward primers containing 20 bp coding the T7 promoter and DreamTaq DNA Polymerase (Thermo Fisher). The PCR products were gel purified using MSB Spin PCRapace (Invitek Molecular) and an *in vitro* transcription was performed using T7 Polymerase (Merck). The remaining DNA was then degraded using DNAse I (Life Technologies). RNA concentration was determined by NanoDrop (Thermo Fisher) and the number of molecules per µL was determined using the online tool EndMemo RNA Copy Number Calculator (https://www.endmemo.com/bio/dnacopynum.php). Primers used can be found in Table S6.

Synchronized worm populations were inoculated with 2.8×10^9^ copies of OrV by pipetting the viral stock on top of the bacterial lawn containing the worms. 700 worms/plate were grown for RT-qPCRs and smiFISH and 1200 worms/plate for the transcriptomic analysis.

#### RNA extractions and RT-qPCRs

Inoculated and control worms were collected at the designated times with PBS-0.05% Tween. Samples were centrifuged for 2 min at 1350 rpm and the supernatant was discarded. Another two wash steps were performed before freezing the samples in liquid nitrogen. 500 µL of Trizol (Invitrogen) were added to the worm pellet and the pellet was disrupted by following 5 cycles of freeze-thawing and 5 cycles of 30 seconds of vortex followed by 30 seconds of rest. 100 µL of chloroform were then added and the tubes were shaken for 15 s and let rest for 2 min. Samples were centrifuged for 15 min at 11,000 g at 4 °C and the top layer containing the RNA was then mixed with the same volume of 100% ethanol. The sample was then loaded into RNA Clean & Concentrator columns (Zymo Research) and the rest of the protocol was followed according to manufacturer instructions.

RT-qPCRs were performed using Power SYBR Green PCR Master Mix (Applied Biosystems) on an ABI StepOne Plus Real-time PCR System (Applied Biosystems). 10 ng of total RNA were loaded and a standard curve was used for OrV quantifications. Primers used for RT-qPCRs can be found in Table S6.

#### Total RNA extraction and preparation for RNA-Seq

Library preparation and Illumina sequencing was done by Novogene Europe (https://www.novogene.com) using a NovaSeq 6000 platform and a Lnc-stranded mRNA-Seq library method, ribosomal RNA depletion and directional library preparation, 150 paired end, and 6 Gb raw data per sample. Novogene checked the quality of the libraries using a Qubit 4 Fluorometer (Thermo Fisher Scientific), qPCR for quantification and Bioanalyzer for size distribution detection.

#### RNA-seq data preprocessing

The quality of the Fastq files was assessed with FASTQC (Andrews, 2010) and MultiQC (Ewels et al., 2016) and preprocessed as paired reads with BBDuk (https://sourceforge.net/projects/bbmap/). Briefly, adapters were eliminated, the first ten 5’ nucleotides of each read were cut, and the 3’ end sequences with average quality below 10 were removed. Processed reads shorter than 60 nucleotides were discarded. (The parameter values were set to: *ktrim* = r, *k* = 31, *mink* = 11, *qtrim* = rl, *trimq* = 10, *maq* = 5, and *forcetrimleft* = 10, and *minlength* = 60).

The resultant processed files were use in the next steps. The N2 Bristol (WBcel235, GCF_000002985.6 assembly) genome was used as reference to the alignment with STAR version 2.7.9a (Dobin et al., 2013), using the run mode “alignReads” and the quantification mode “GeneCounts”. Quantification results were used for the transcriptome analysis.

#### Differential expression analysis and functional enrichment analysis

DESEq2 v1.38.3 (Love et al., 2014) was used for the DEA using test = “LRT” (complete design = condition + time + condition:time, reduced design = condition + time). The DEGs by timepoint were extracted with test = “Wald” on each result (time). DEGs were considered significant with Benjamini and Hochberg adjusted *P* ≤ 0.05 values. In most heatmaps asterisks indicate significance level by timepoint.

The FEAs were performed using the WormEnrichr site with the enrichr package v 3.2 (Kuleshov et al., 2016). Only terms with adjusted *P* ≤ 0.05 were considered.

#### Correlation with virus accumulation

A Pearson’s correlation between the differential expression of the DEGs (log2-fold change by timepoint) and the virus accumulation (percentage of viral reads in sample) was performed with cor.test function from stats package v.4.2.1. Only significant DEGs were evaluated. Genes with an *r* ≥ 0.8 were considered correlated and those with an *r* ≤ −0.8 were considered to be anticorrelated.

#### PPI Network functional enrichment and visualization

The PPI network of the specific genes in the OrV response was created with Cytoscape v3.10.0 (Shannon et al., 2003) selecting the option “Physical subnetwork”. The enrichment and visualization of the categories was performed with the STRING plugin.

#### smiFISH

The smiFISH protocol was adapted from previous studies (Tsanov et al., 2016; Parker et al., 2021). Inoculated and control worms were washed with PBS 0.05% Tween and centrifuged for 2 min at 1350 rpm four times discarding the supernatants. 800 µL of Bouins fixation mix (400 µL Bouins Fix, 400 µL methanol, 10 µL β-mercaptoethanol) were added and the sample incubated at room temperature for 30 min in a rotatory shaker. Samples were then frozen in liquid nitrogen and kept at −80 °C overnight. The following day samples were gently shaken at 4 °C for 30 min and washed 4 times with borate Triton solution (20 mM H3BO3, 10 mM NaOH, 0.5% Triton) and 5 times with borate Triton β-mercaptoethanol solution (20 mM H3BO3, 10 mM NaOH, 0.5% Triton, 2% β-mercaptoethanol), leaving the borate Triton β-mercaptoethanol incubate for 1 h between each wash in the last 3 washes. The sample was then incubated for 5 min with wash buffer A (1 mL Stellaris RNA FISH Wash Buffer A, 1 mL deionized formamide and 3 mL H2O).

Forty probes against OrV were designed using Oligostan (Tsanov et al., 2016), 22 against RNA1 and 18 against RNA2. The sequences of the probes can be found in Table S1. The probes were annealed as follows: 2 µL 0.83 µM probe set, 1 µL 50 µM FLAP-label, 1 µL NEB3 and 6 µL DEPC H2O were incubated at 85 °C for 3 min, 65 °C for 3 min and 25 °C for 5 min. The FLAP labels contained CAL Fluor 610 or Quasar 670 modifications at both the 5’ and 3’ ends of the following sequence: 5’ AATGCATGTCGACGAGGTCCGAGTGT (Biosearch Technologies).

One µL of annealed probe solution targeting RNA1 and 1 µL of annealed probe solution targeting RNA2 was then mixed with 98 µL of hybridization buffer (100 µL Stellaris smiFISH hybridization buffer, 25 µL Deionized formamide). 100 µL of hybridization buffer containing the smiFISH probes were then added to the samples and the samples were incubated overnight at 37 °C in a rotary shaker. The following day, samples were washed with wash buffer A2 (1 mL, Stellaris RNA FISH Wash Buffer A, 4 mL H2O), incubated for 30 min in wash buffer A2 at 37 °C, and then incubated for 30 min in Wash buffer A2 containing 25 ng of DAPI at 37 °C. Samples were then incubated 5 minutes in Stellaris RNA FISH Wash Buffer B, centrifuged and resuspended in a small volume of Stellaris RNA FISH Wash Buffer B. 0.1 ng of DAPI were added to each sample and samples were mounted using N-propyl gallate mounting medium. Samples were imaged using Leica DMi8 microscope with Leica DFC9000 GTC sCMOS camera and objective HC PL APO 40x/0,95 CORR PH2. Images were analyzed and processed using ImageJ(FIJI) (Schindelin et al., 2012).

smiFISH signal was used to determine the number of infected cells in each worm, the percentage of worms with RNA2 vs RNA1 and the localization of the signal.

#### Percentage of infected worms

In order to determine the percentage of infected worms the presence or absence of smiFISH, *pals5p::GFP* or *lys-3p::GFP* signal was used to determine the infection status of each worm. From the total number of worms analyzed, the percentage of infected worms was calculated.

#### Clustering of gene expression profiles

All the significant genes (adjusted *P* ≤ 0.05) identified with the LRT were clustered using the *hclust* method from stats package v.4.2.1. A co-occurrence matrix was generated from the results through cutting the tree from 2 to 100 groups (cutree, *k* = 2:100). Only the genes that belonged to the same cluster in at least the 50% of the *k* were treated as connected for the adjacency matrix. To identify the modules, we used the method *clusters* from the igraph package v1.5.0 (Csardi and Nepusz, 2006), keeping those formed by a minimum of five genes.

#### Comparison with other biotic stresses

The Microbes datasets from WormExp v2.0 (Yang et al., 2016) was used for the comparison with other biotic stresses with the limitation that only experiments performed on wild-type N2 strain were considered. Data from two additional experiments in which N2 strain was used and at least three biological replicates were done was also included (Chen et al., 2017; Osman et al., 2018). A total of 55 experiments were used. Our significant DEGs by time point (adjusted *P* ≤ 0.0499) were used in the comparison. All pathogens were classified into the following categories: virus, Gram− bacteria, Gram+ bacteria and fungi. A gene was considered common when identified as DEGs (adjusted *P* ≤ 0.05) in at least one experiment of each category.

#### Lipids metabolism, antimicrobial effectors, IPR and *zip-1* dependent genes

*C. elegans* genes related to the lipid metabolism (GO0006629) where downloaded from Wormbase database. IPR genes were extracted from Table S8 (Reddy et al., 2019) and ZIP-1 dependent genes from Table S3 (Lažetić et al., 2022).

#### Code and machine specifications

All the code generated for this study is accessible in the repository https://git.csic.es/sfelenalab/OrV_progression. The more intensive computations (preprocessing of raw fastqs and alignments) were performed on the HPC cluster Garnatxa at I2SysBio (CSIC-UV). The rest of the analysis (mainly in R version 4.2.1, 2022-06-23), were ran in an iMac (2020) with a 3.3 GHz Intel Core i5 of 6 cores, with 16 GB (2667 MHz DDR4) RAM.

#### *gst-35* CRISPR knock-out

The *gst-35(esv1)* deletion was generated by imprecise repair of a CRISPR/Cas9-induced DNA double-strand break and selected using the *dpy-10* co-CRISPR approach (Arribere et al., 2014). The fusion was generated using plasmid-based expression of Cas9 and sgRNAs. Two pairs of sgRNA plasmids were used to target the 5’ and 3’ ends of the *gst-35* open reading frame and generated by ligation of annealed oligo pairs into the pU6::sgRNA expression vector pJJR50 (Addgene) as previously described (Waaijers et al., 2016). The sgRNA targeting the *dpy-10* locus (Arribere et al., 2014) was cloned into the pU6::sgRNA expression vector pJJR50 as previously described (Waaijers et al., 2016). The injection mix was prepared in MilliQ H2O and contained 60 ng/mL Peft-3::Cas9 (Addgene) and 45 ng/mL each sgRNA plasmid. Young adult hermaphrodites were injected in the germline. For selection of edited genomes, injected animals were transferred to individual culture plates, incubated for 3 to 4 d at 20 °C, and 96 non-transgenic F1 animals (WT, Dpy, or Rol) from two plates containing high numbers of Dpy and Rol animals were selected and transferred to individual NGM plates. After laying eggs, F1 animals were lysed and genotyped by PCR using two primers flanking the *gst-35* open reading frame. Sanger sequencing was used to determine the precise molecular lesion (Eurofins Genomics). The sequences targeted and the primers used can be found in Table S6.

## SUPPLEMENTAL INFORMATION TITLE AND LEGENDS

Figure S1. OrV positive worms in wild-type and sensitive background JU1580 strains Percentage of *pals-5p*::GFP positive worms in the N2 background and *lys-3p*::GFP positive worms in the sensitive JU1580 background. Worms were inoculated with 2.25×10^9^ copies of OrV and analyzed at 48 hpi. *n* = 106 worms for ERT54 and 91 for JU1580, within 5 replicates.

Figure S2. Functional enrichment by time point Up (cyan) and down (magenta) -regulated DEGs where analyzed separately. Only significant terms are shown (adjusted *P* ≤ 0.05).

(A) GO:BP.

(B) GO:MF.

(C) KEGG pathways.

Figure S3. Ubiquitous genes

(A) Expression profile of the 29 genes common to the exponential viral replication and host-pathogen conflict stages. All the genes present an up-regulated profile. The asterisks indicate the significance level at each time point: \**P* ≤ 0.05, \*\**P* ≤ 0.01 and \*\*\**P* ≤ 0.001.

(B) Genes description.

Figure S4. Representative profiles of the clusters All the significant DEGs (LRT analysis) where clustered as indicated in Material and Methods. The mean *z*-score (±1 SD) of the control (black) and infected (purple) groups are represented. Only clusters with at least 5 genes are shown.

Figure S5. Biotic stress comparison analysis Pathogens included in the comparison. Expression profile of the OrV-specific genes that form part of the PPI network shown in Fig. 6C. OrV-regulated coding and non-coding sequence types.

Figure S6. Differential expression of the DEGs related with the Intracellular Pathogen Response in our time series experiments and comparison with other biotic stresses

(A) Differential expression of IPR genes affected in our time series data. Asterisks indicate significance by timepoint (adjusted significance levels: \**P* ≤ 0.05, \*\**P* ≤ 0.01 and \*\*\**P* ≤ 0.001).

(B) Differential expression of the genes in A upon various biotic stresses. Cyan indicates upregulation and magenta indicates downregulation (adjusted *P* ≤ 0.05).

Figure S7. Differential expression of antimicrobial effectors identified as DEGs in our time series experiments and comparison with other biotic stresses

(A)Differential expression of antimicrobial effector genes affected in our time series data. Asterisks indicate significance by timepoint (adjusted significance levels: \**P* ≤ 0.05, \*\**P* ≤ 0.01 and \*\*\**P* ≤ 0.001).

(B) Differential expression of genes in A upon various biotic stresses. Cyan indicates upregulation and magenta indicates downregulation (adjusted *P* ≤ 0.05).

Figure S8. Differential expression of DEGs related with the lipid metabolism in our time series experiments and comparison with other biotic stresses

(A) Differential expression of lipid metabolism related genes affected in our time series data. Asterisks indicate significance by timepoint (adjusted significance levels: \**P* ≤ 0.05, \*\**P* ≤ 0.01 and \*\*\**P* ≤ 0.001).

(B) Differential expression of the genes in A upon various biotic stresses (all performed on the *wild-type*). Cyan indicates upregulation and magenta indicates downregulation (adjusted *P* ≤ 0.05).

Figure S9. Expression of interesting DEGs after OrV exposure

(A) Expression profiles of selected DEGs.

(B) Schematic representation of *gst-35(esv1)* deletion.

(C) *gst-35* CRISPR design.

(D) *gst-35(esv1)* allele.

(E) GST-35 protein and *esv1* deletion with horizontal bars indicating the conserved amino acids.

## EXCEL TABLE TITLES AND LEGENDS

Table S1. DEA significant results.

Table S2. Functional enrichment complete results.

Table S3. Clusters LRT DEGs.

Table S4. Biotic stresses experiments comparison.

Table S5. Overlap between OrV-specific responding genes and DEGs of *C. briggsae* upon Santeuli virus infection.

Table S6. Primers

Table S7. Key resource table

