## Supplementary figures for "Story of an infection: viral dynamics and host responses in the *Caenorhabditis elegans*-Orsay virus pathosystem": Supplementary figures.pptx

### Slide 1
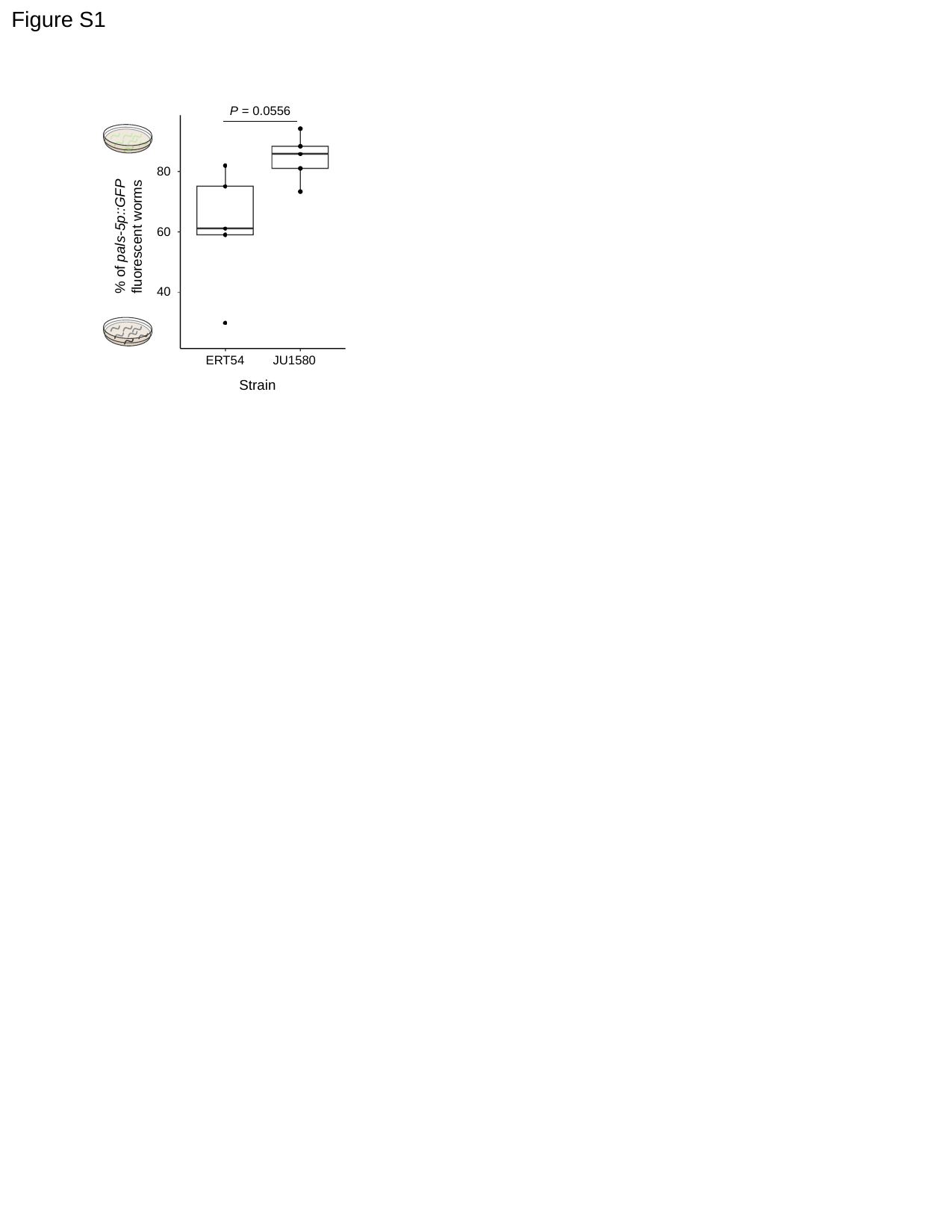

Figure S1
P = 0.0556
80
% of pals-5p::GFP
fluorescent worms
60
40
ERT54
JU1580
Strain

### Slide 2
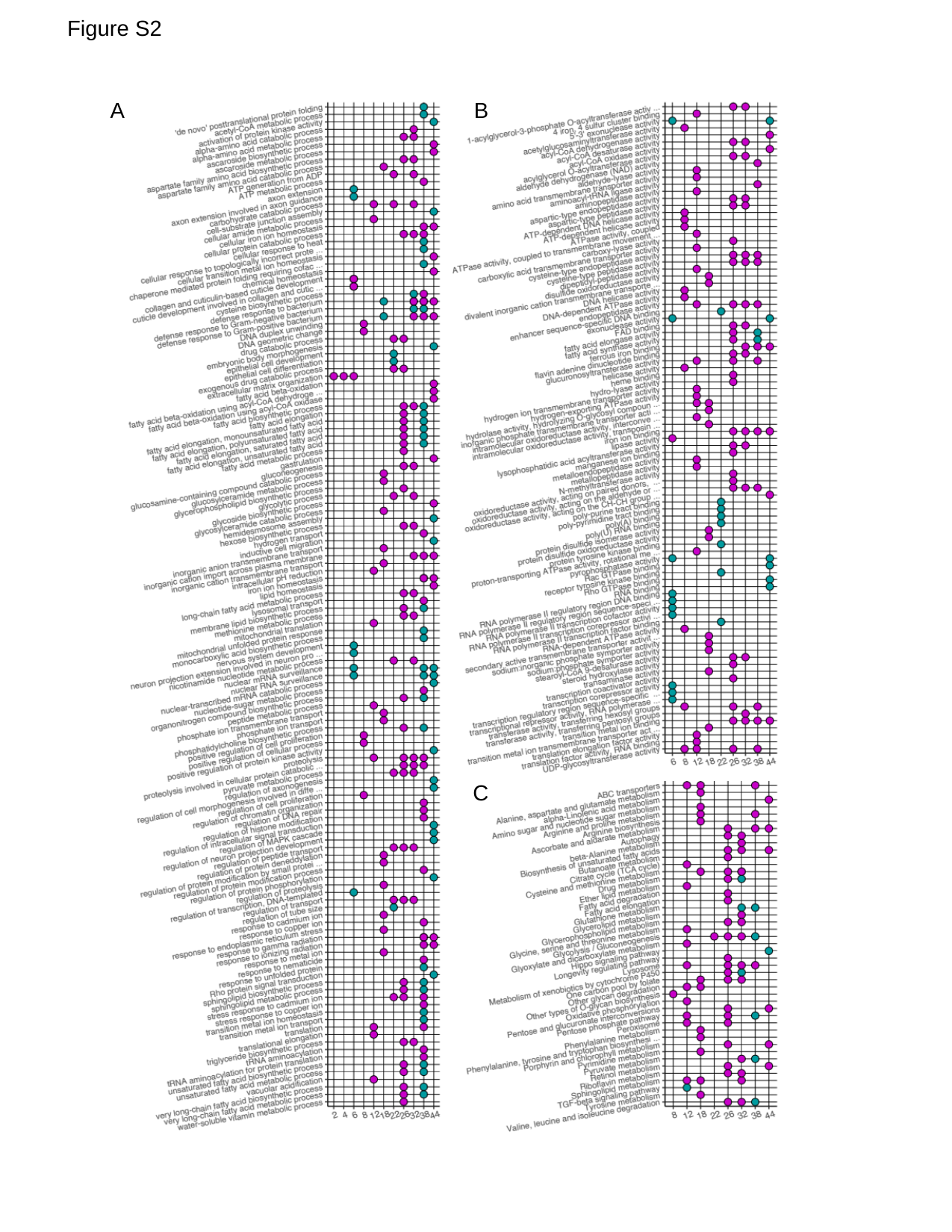

Figure S2
B
A
C

### Slide 3
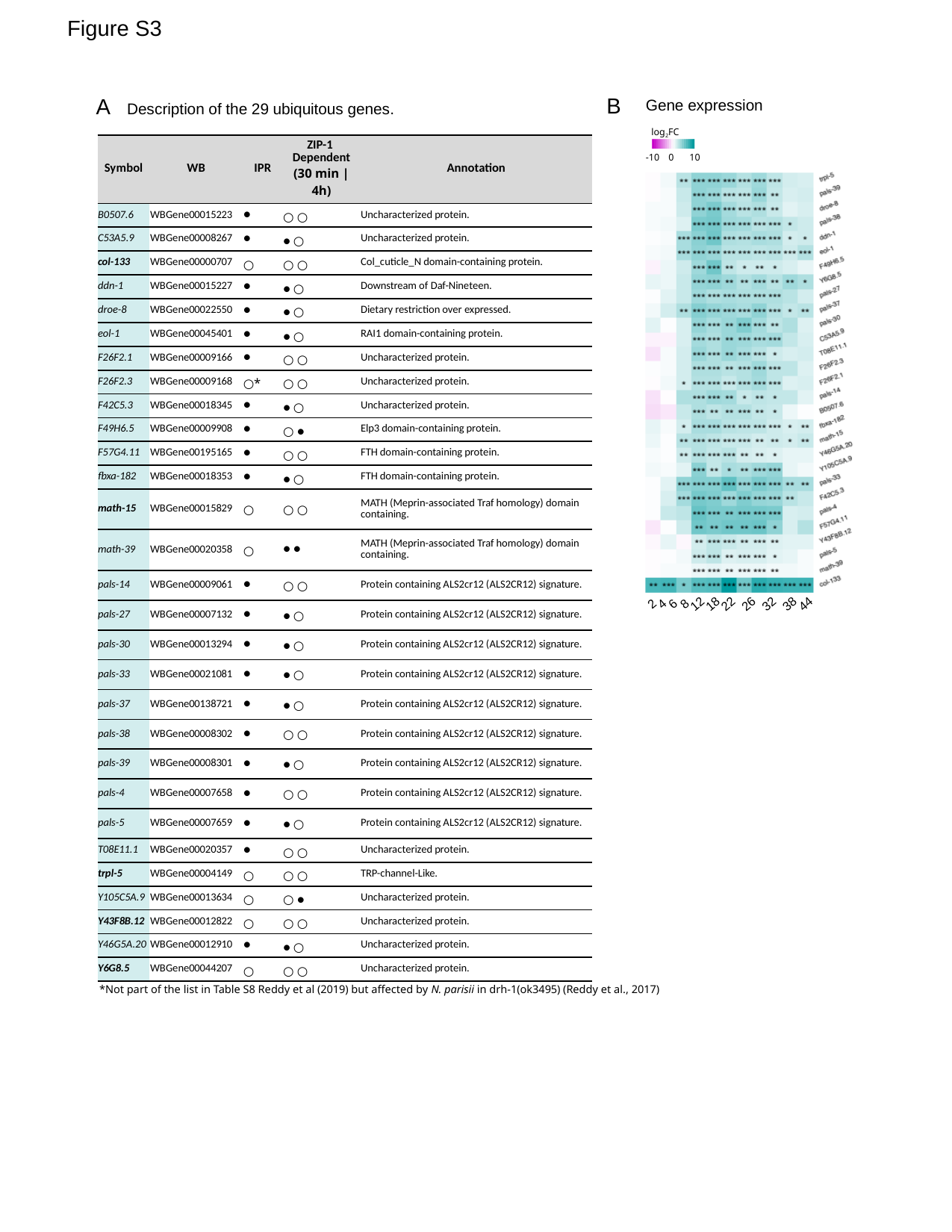

>Mas correlación carga viral
Genes desincronizados
Male L4
Dauer
>Enriquecimiento funcional (next slide)
#### Desarollo – heatmaps / experimental?
Figure S3
B
A
Gene expression
log_2_FC
-10
0
10
2
4
6
8
12
18
22
26
32
38
44
Description of the 29 ubiquitous genes.
| Symbol | WB | IPR | ZIP-1 Dependent (30 min | 4h) | Annotation |
| --- | --- | --- | --- | --- |
| B0507.6 | WBGene00015223 | ● | ○ ○ | Uncharacterized protein. |
| C53A5.9 | WBGene00008267 | ● | ● ○ | Uncharacterized protein. |
| col-133 | WBGene00000707 | ○ | ○ ○ | Col\_cuticle\_N domain-containing protein. |
| ddn-1 | WBGene00015227 | ● | ● ○ | Downstream of Daf-Nineteen. |
| droe-8 | WBGene00022550 | ● | ● ○ | Dietary restriction over expressed. |
| eol-1 | WBGene00045401 | ● | ● ○ | RAI1 domain-containing protein. |
| F26F2.1 | WBGene00009166 | ● | ○ ○ | Uncharacterized protein. |
| F26F2.3 | WBGene00009168 | ○\* | ○ ○ | Uncharacterized protein. |
| F42C5.3 | WBGene00018345 | ● | ● ○ | Uncharacterized protein. |
| F49H6.5 | WBGene00009908 | ● | ○ ● | Elp3 domain-containing protein. |
| F57G4.11 | WBGene00195165 | ● | ○ ○ | FTH domain-containing protein. |
| fbxa-182 | WBGene00018353 | ● | ● ○ | FTH domain-containing protein. |
| math-15 | WBGene00015829 | ○ | ○ ○ | MATH (Meprin-associated Traf homology) domain containing. |
| math-39 | WBGene00020358 | ○ | ● ● | MATH (Meprin-associated Traf homology) domain containing. |
| pals-14 | WBGene00009061 | ● | ○ ○ | Protein containing ALS2cr12 (ALS2CR12) signature. |
| pals-27 | WBGene00007132 | ● | ● ○ | Protein containing ALS2cr12 (ALS2CR12) signature. |
| pals-30 | WBGene00013294 | ● | ● ○ | Protein containing ALS2cr12 (ALS2CR12) signature. |
| pals-33 | WBGene00021081 | ● | ● ○ | Protein containing ALS2cr12 (ALS2CR12) signature. |
| pals-37 | WBGene00138721 | ● | ● ○ | Protein containing ALS2cr12 (ALS2CR12) signature. |
| pals-38 | WBGene00008302 | ● | ○ ○ | Protein containing ALS2cr12 (ALS2CR12) signature. |
| pals-39 | WBGene00008301 | ● | ● ○ | Protein containing ALS2cr12 (ALS2CR12) signature. |
| pals-4 | WBGene00007658 | ● | ○ ○ | Protein containing ALS2cr12 (ALS2CR12) signature. |
| pals-5 | WBGene00007659 | ● | ● ○ | Protein containing ALS2cr12 (ALS2CR12) signature. |
| T08E11.1 | WBGene00020357 | ● | ○ ○ | Uncharacterized protein. |
| trpl-5 | WBGene00004149 | ○ | ○ ○ | TRP-channel-Like. |
| Y105C5A.9 | WBGene00013634 | ○ | ○ ● | Uncharacterized protein. |
| Y43F8B.12 | WBGene00012822 | ○ | ○ ○ | Uncharacterized protein. |
| Y46G5A.20 | WBGene00012910 | ● | ● ○ | Uncharacterized protein. |
| Y6G8.5 | WBGene00044207 | ○ | ○ ○ | Uncharacterized protein. |
*Not part of the list in Table S8 Reddy et al (2019) but affected by N. parisii in drh-1(ok3495) (Reddy et al., 2017)

### Slide 4
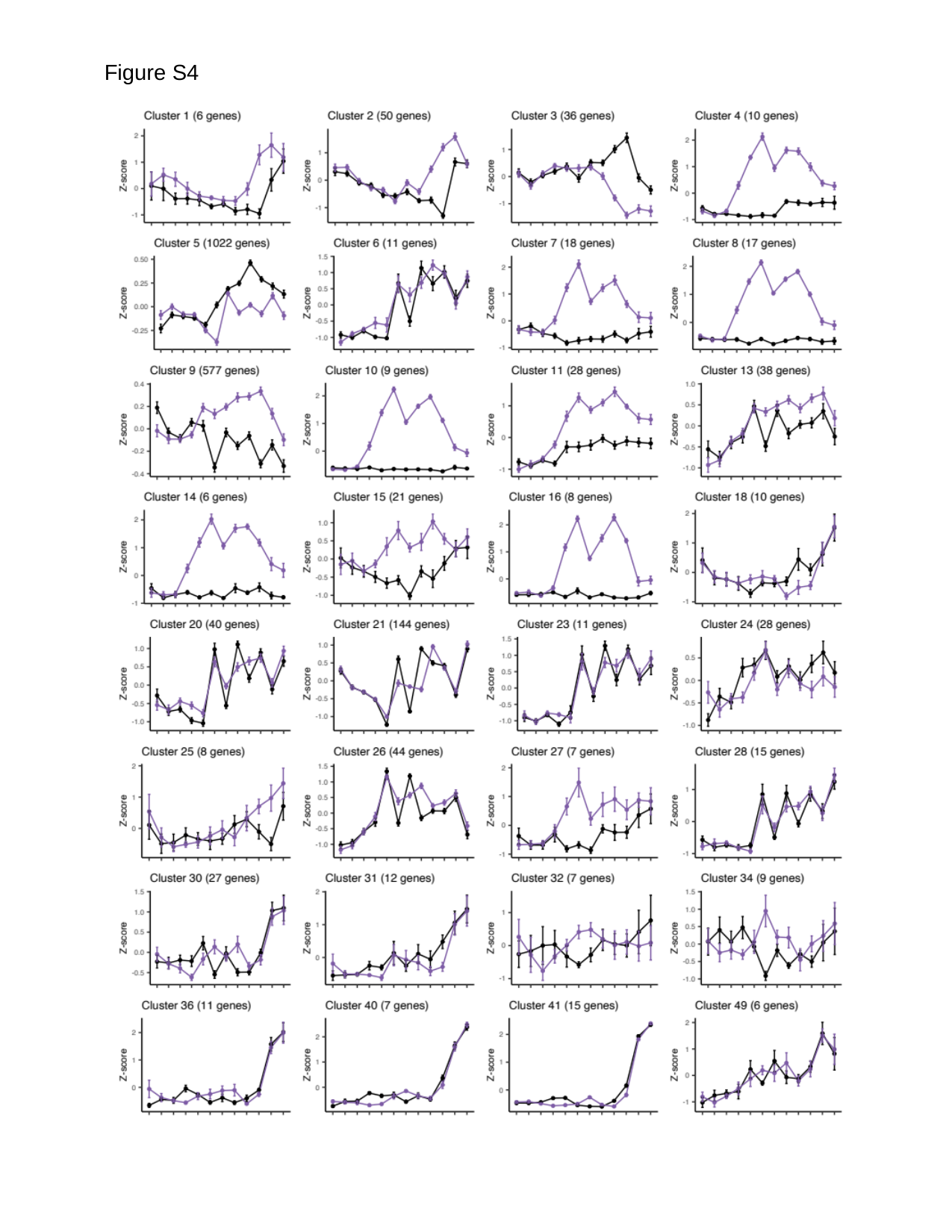

Figure S4

### Slide 5
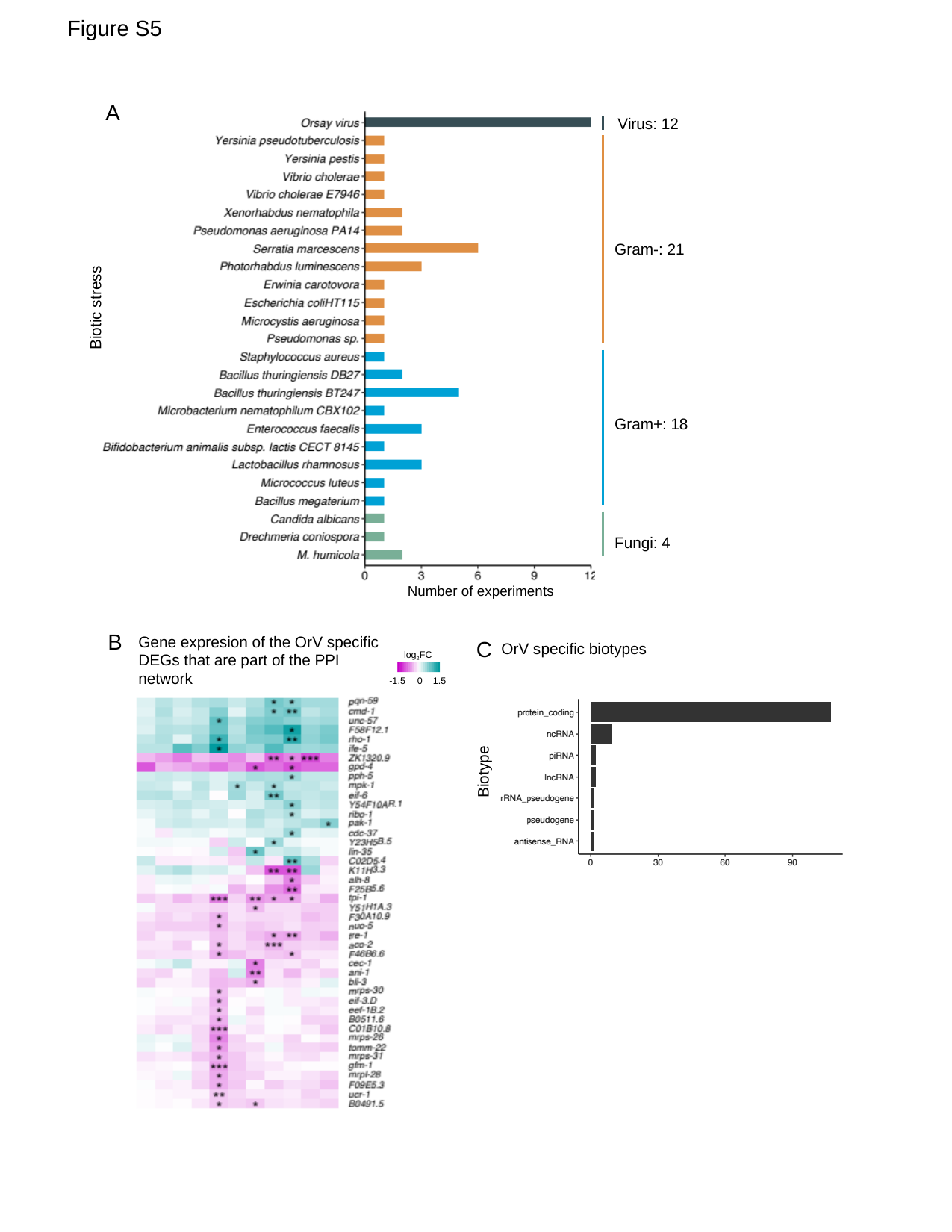

Figure S5
A
Virus: 12
Gram-: 21
Biotic stress
Gram+: 18
Fungi: 4
Number of experiments
B
Gene expresion of the OrV specific DEGs that are part of the PPI network
C
OrV specific biotypes
log_2_FC
-1.5
0
1.5
Biotype

### Slide 6
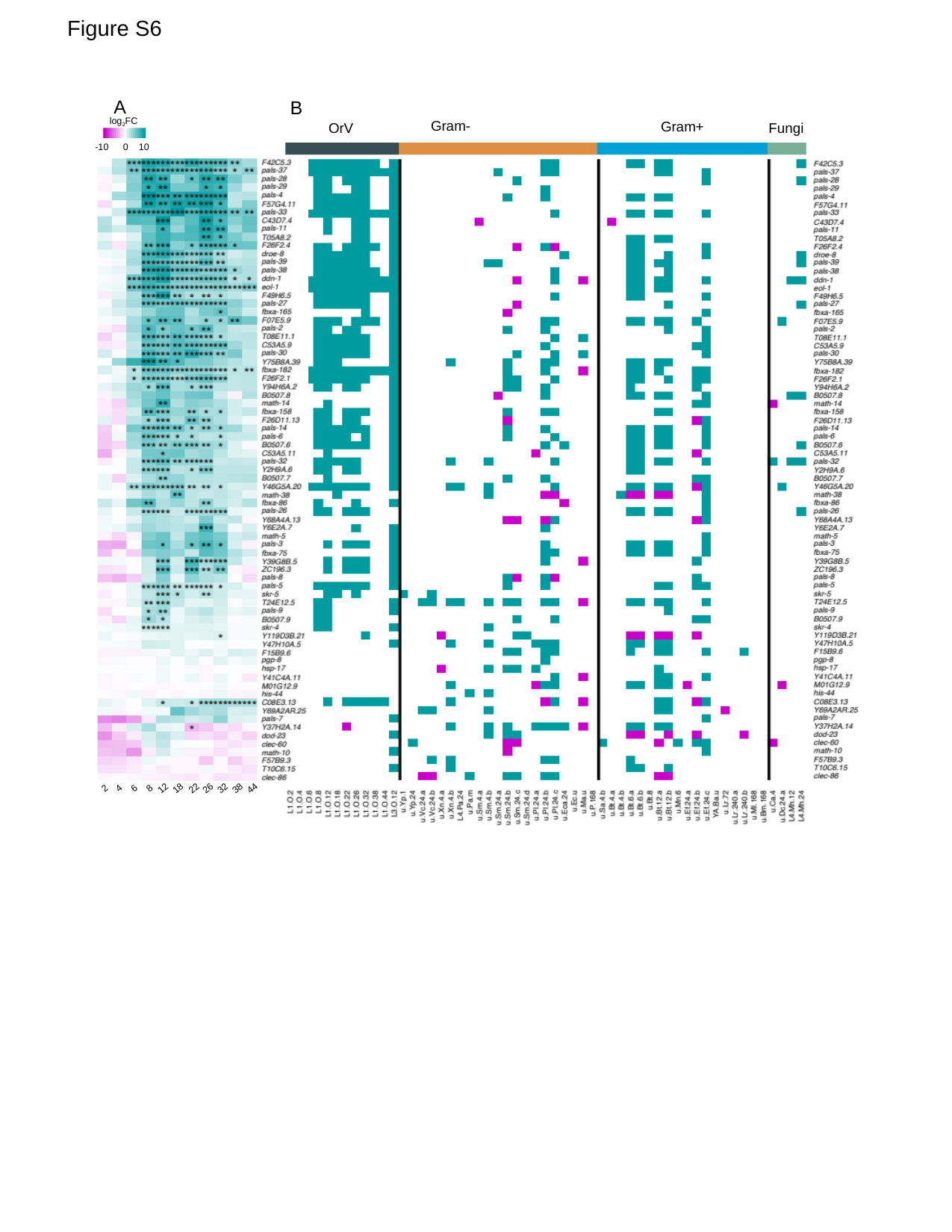

Figure S6
A
B
log_2_FC
-10
0
10
Gram-
Gram+
OrV
Fungi
2
4
6
8
12
18
22
26
32
38
44
2
4
6
8
12
18
22
26
32
38
44

### Slide 7
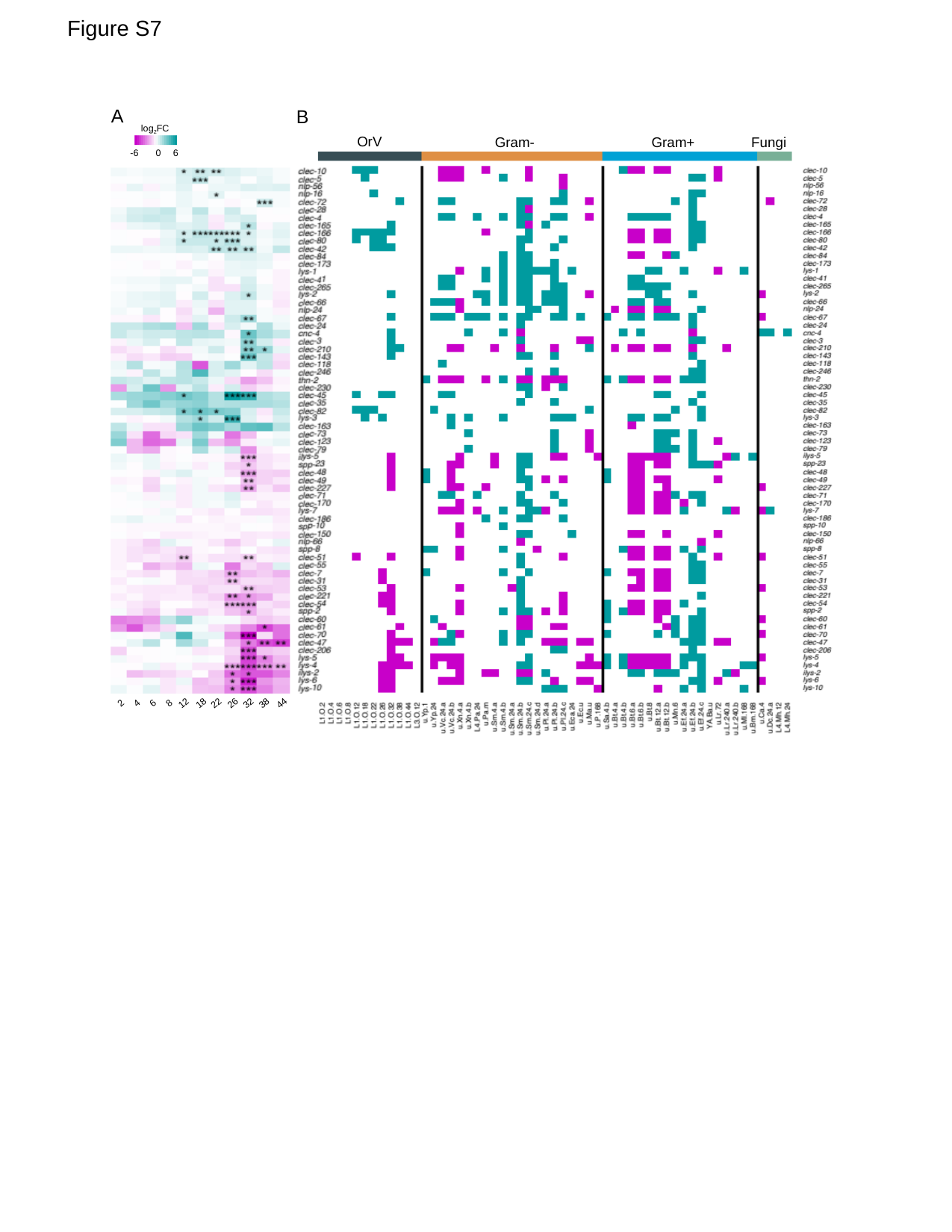

Antimicrobial effectors
Figure S7
A
B
log_2_FC
-6
0
6
OrV
Gram-
Gram+
Fungi
2
4
6
8
12
18
22
26
32
38
44

### Slide 8
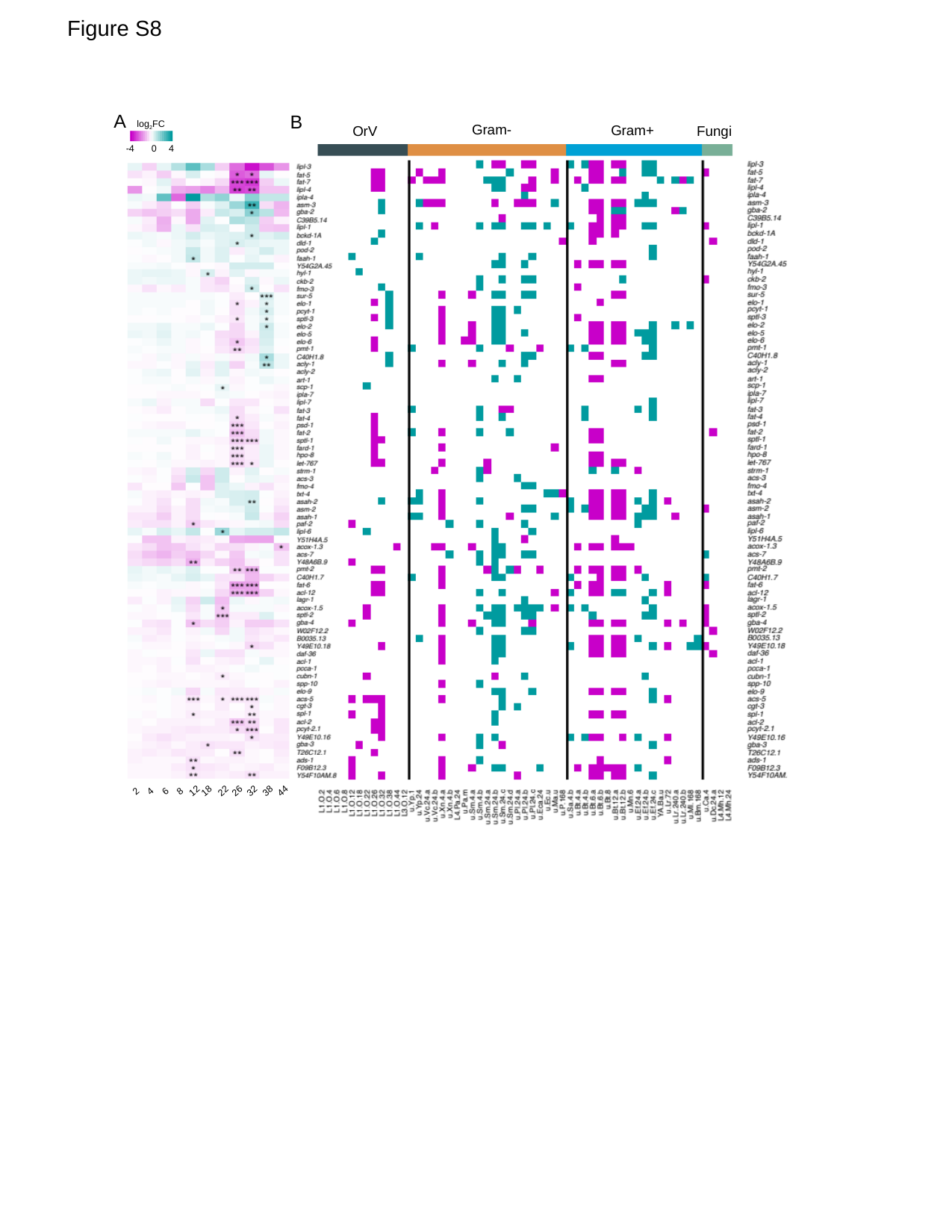

Figure S8
A
B
log_2_FC
-4
0
4
Gram-
Gram+
Fungi
OrV
2
4
6
8
12
18
22
26
32
38
44
2
4
6
8
12
18
22
26
32
38
44

### Slide 9
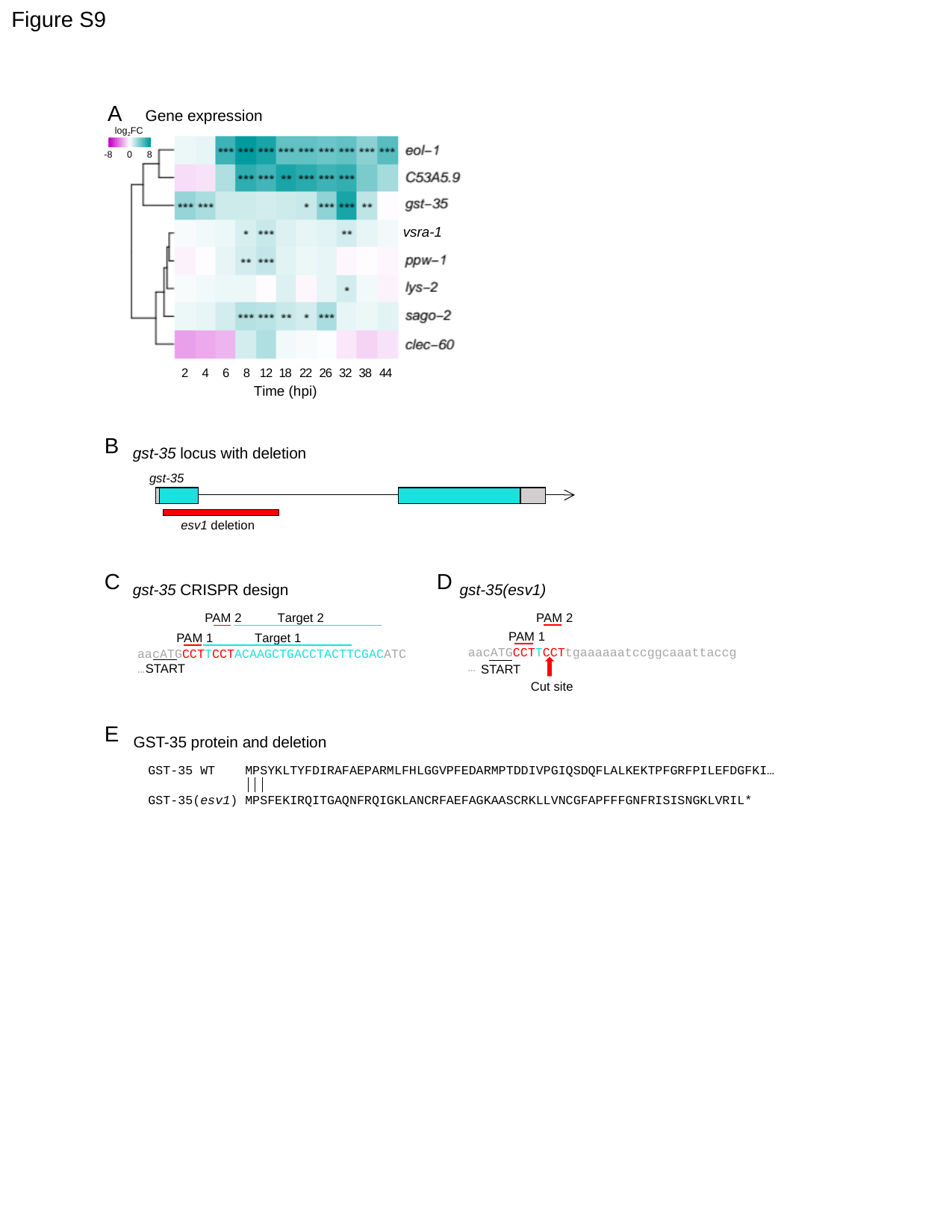

Figure S9
A
Gene expression
log_2_FC
-8
0
8
vsra-1
2
4
6
8
12
18
22
26
32
38
44
Time (hpi)
B
gst-35 locus with deletion
gst-35
esv1 deletion
C
D
gst-35 CRISPR design
gst-35(esv1)
PAM 2
PAM 2
Target 2
PAM 1
PAM 1
Target 1
aacATGCCTTCCTtgaaaaaatccggcaaattaccg…
aacATGCCTTCCTACAAGCTGACCTACTTCGACATC…
START
START
Cut site
E
GST-35 protein and deletion
GST-35 WT MPSYKLTYFDIRAFAEPARMLFHLGGVPFEDARMPTDDIVPGIQSDQFLALKEKTPFGRFPILEFDGFKI…
GST-35(esv1) MPSFEKIRQITGAQNFRQIGKLANCRFAEFAGKAASCRKLLVNCGFAPFFFGNFRISISNGKLVRIL*
