## Supplementary Tables for "Story of an infection: viral dynamics and host responses in the *Caenorhabditis elegans*-Orsay virus pathosystem": Supplementary Table S7_key resources.docx

| **0/0/00 0:00:00REAGENT or RESOURCE** | **SOURCE** | **IDENTIFIER** |
| --- | --- | --- |
| **Deposited data** | | |
| **Bacterial and virus strains** | | |
| *Escherichia coli* OP50 | Caenorhabditis Genetics Center (CGC) | RRID: WB-STRAIN:OP50 |
| Orsay virus | Marie-Anne Félix's lab (Félix et al., 2011) | NCBI:txid977912 |
| **Experimental models: Organisms/strains** | | |
| *C. elegans*: Strain: JU2624: *[myo-2::mcherry::unc54; lys-3p::eGFP::tbb-2]* *IV* in JU1580 background | Marie-Anne Félix's lab | N/A |
| *C. elegans*: Strain: SFE2: *drh-1(ok3495)IV; mjIs228[myo-2::mcherry::unc54; lys-3p::eGFP::tbb-2]?* | This study | N/A |
| *C. elegans*: Strain: ERT54: *jyIs8[pals-5p::GFP; myo-2p::mCherry] X* | Emily Troemel's lab | WB Strain: ERT54 |
| *C. elegans*: Strain: RB2519: *drh-1(ok3495)IV* | CGC | WB Strain: RB2519 |
| *C. elegans*: Strain: ERT288: *pals-22(jy3) III; jyIs8[pals-5p::GFP, myo-2::mCherry] X* | Emily Troemel's lab | WB Strain: ERT288 |
| *C. elegans*: Strain: NL2550: *ppw-1(pk2505) I* | CGC | WB Strain: NL2250 |
| *C. elegans*: Strain: WM154: *sago-2(tm894) I* | CGC | WB Strain: WM154 |
| *C. elegans*: Strain: WM153: *C04F12.1(tm1637) I* | CGC | WB Strain: WM153 |
| *C. elegans*: Strain: SFE9: *lys-2(tm2398); jyIs8[pals-5p::GFP + myo-2p::mCherry]X* | This study | N/A |
| *C. elegans*: Strain: CB6734: *clec-60(tm2319) II* | CGC | WB Strain: CB6734 |
| *C. elegans*: Strain: SFE10: *eol-1(tm6609); jyIs8[pals-5p::GFP + myo-2p::mCherry]X* | This study | N/A |
| *C. elegans*: Strain: SFE7: *C53A5.9(tm10539); jyIs8[pals-5p::GFP + myo-2p::mCherry]X* | This study | N/A |
| *C. elegans*: Strain: SFE11: *gst-35(esv1) II* | This study | N/A |
| **Oligonucleotides** | | |
| See Table S6 for oligonucleotides |  |  |
| **Software and algorithms** | | |
| FastQC v0.12.1 | Andrews^1^ | <https://www.bioinformatics.babraham.ac.uk/projects/fastqc/> |
| MultiQC v1.12 | Ewels et al.^2^ | <https://multiqc.info> |
| bbduk.sh v39.01 |  | <https://sourceforge.net/projects/bbmap/>  https://jgi.doe.gov/data-and-tools/software-tools/bbtools/bb-tools-user-guide/bbduk-guide/ |
| STAR v.2.7.9a | Dobin et al.^3^ | <https://github.com/alexdobin/STAR> |
| R version 4.2.1 (2022-06-23) | R Core Team^4^ | <https://www.r-project.org> |
| RStudio 2022.02.3+492 "Prairie Trillium" |  | <https://posit.co/products/open-source/rstudio/> |
| R packages and versions listed in sessionInfo | This study | <https://git.csic.es/sfelenalab/OrV_progression/blob/main/sessionInfo.txt> |
| **Other** | | |
| Scripts created to perform the bioinformatic analysis: from raw fastq files to the plots | This study | <https://git.csic.es/sfelenalab/>OrV_progression/tree/main |

4.	R Core Team. *R: A Language and Environment for Statistical Computing*. (R Foundation for Statistical Computing, 2022).
